# An unconventional autophagic pathway that inhibits ATP secretion during apoptotic cell death

**DOI:** 10.1101/2024.01.21.576513

**Authors:** Elena Terraza-Silvestre, Raquel Villamuera, Julia Bandera-Linero, Michal Letek, Cristina Ramón-Barros, Clara Moyano-Jimeno, Daniel Oña-Sánchez, Felipe X. Pimentel-Muiños

## Abstract

Mobilization of Damage-Associated Molecular Patterns (DAMPs) determines the immunogenic properties of apoptosis, but the mechanisms that control DAMP exposure are still unclear. Here we describe an unconventional autophagic pathway that inhibits the release of ATP, a critical DAMP in immunogenic apoptosis, from dying cells. Mitochondrial BAK activated by BH3-only molecules interacts with prohibitins and stomatin-1 through its latch domain, indicating the existence of an interactome specifically assembled by unfolded BAK. This complex engages the WD40 domain of the critical autophagic effector ATG16L1 to induce unconventional autophagy, and the resulting LC3-positive vesicles contain ATP. Functional interference with the pathway increases ATP release during cell death, reduces ATP levels remaining in the apoptotic bodies, and improves phagocyte activation. These results reveal that an unconventional component of the autophagic burst that often accompanies apoptosis sequesters intracellular ATP to prevent its release, thus favoring the immunosilent nature of apoptotic cell death.

## INTRODUCTION

The BCL2 protein family plays essential roles in the intrinsic apoptotic pathway, which has important developmental and homeostatic functions(*1*)(*2*)(*3*). The intrinsic route is initiated by activation of BH3-only proteins, a BCL2-family subgroup including just one of the prototypical <u>B</u>CL2-<u>H</u>omology (BH) domains. Induced BH3-only effectors activate the multidomain molecules BAK and BAX to cause their partial unfolding and multimerization, thus opening high-conductance pores on the outer mitochondrial membrane (OMM) for cytochrome c release and caspase activation(*4*)(*5*)(*6*)(*7*).

Intrinsic apoptosis in homeostatic conditions is normally immunosilent, in such a way that specialized phagocytes remove the cellular remains without triggering an immune response(*8*). However, in reaction to certain drugs and/or conditions, apoptosis can be strongly immunogenic, thus sparking an adaptive immune reaction against neoantigens that may be present in the dying cells(*9*). Immunogenic cell death favors long-lasting tumor remission by promoting anti-tumor immunity, and therefore plays a critical role in cancer therapy(*10*)(*11*)(*12*). Induction of immunogenic cell death requires the release of adjuvant factors by the dying cells that stimulate the pro-inflammatory properties of the phagocytes that eliminate the cellular remains(*13*). Such factors are known as Damage-Associated Molecular Patterns (DAMPs), and include exposure of calreticulin at the surface of the apoptotic cell, release of nuclear HMGB1 and ATP secretion(*14*)(*15*). Secreted ATP acts through purinergic receptors as a chemotactic agent for phagocyte recruitment and as a phagocyte activation factor for pro-inflammatory cytokine secretion(*16*), and plays a central role in immunogenic cell death. The mechanisms that regulate DAMP emission, and how they are connected to the apoptotic machinery, are still unclear, but autophagy has been involved in both HMGB1 and ATP secretion(*17*)(*18*)(*19*)(*20*)(*21*), with a particularly relevant function described for the latter in anti-cancer immunity induced by chemotherapy(*18*).

Autophagy, a catabolic process that degrades cytoplasmic components through the action of <u>A</u>u<u>T</u>opha<u>G</u>y-related proteins (ATGs)(*22*), is often induced during cell death, suggesting that its degradative activity may contribute to the cell’s suicide. However, autophagy does not usually play an execution role, but rather it accompanies apoptosis with both contributory or inhibitory roles previously described(*23*). While the canonical autophagic pathway is well characterized, different ATGs participate in a variety of atypical autophagic activities that are mechanistically unconventional and/or unrelated to the degradative purpose of autophagy(*24*)(*25*)(*26*)(*27*). These processes are collectively known as unconventional autophagy and serve a variety of purposes, like elimination of invading agents(*28*), detection of lysosomal dysfunction(*29*), protein secretion(*30*)(*31*)(*32*) and trafficking(*33*)(*34*)(*35*), inflammatory control(*36*)(*37*) or receptor signaling(*38*). Notably, some of these activities involve lipidation of the autophagic marker LC3 in single-membrane vesicles unrelated to canonical double-membrane autophagosomes(*28*)(*39*)(*35*)(*33*)(*34*). This atypical LC3 behavior often requires the C-terminal domain of the critical effector ATG16L1, a region of the molecule that includes seven WD40-type repeats (WD40 domain, WDD) and is unnecessary for canonical autophagy(*33*)(*40*)(*41*)(*39*). Given this functional plurality, detection of a certain autophagic activity may reflect a variety of underlying mechanisms and physiological roles. The exact nature and functional implications of the autophagic burst that often accompanies apoptosis are issues that remain to be fully elucidated.

Here we show that mitochondrial BAK activated by BH3-only molecules induces an unconventional autophagic response that suppresses the release of ATP during apoptotic cell death by sequestering it into atypical autophagic vesicles, thus limiting the immunogenic potential of the dying cells. We provide evidence of a novel mitochondrial complex assembled by activated BAK to induce this activity.

## RESULTS

### BAK mediates autophagy activated during apoptotic cell death

To study how autophagy is modulated during apoptosis we retrovirally expressed BH3-only molecules in mouse embryonic fibroblasts (MEFs) as a way to dissect the molecular consequences of exciting a single pathway and avoid the noise caused by chemicals that often stimulate a variety of stress signaling routes. We first transduced different BH3-only members into wild-type (WT) MEFs in the presence of the caspase inhibitor zVAD.fmk to delay cell death. Out of the eight BH3-only molecules tested, only BIM, tBID and PUMA induced robust accumulation of LC3-II, a common autophagic reporter (Figure S1A). Expression of the positive effectors did not activate LC3 in *Bak/Bax*-double knock-out MEFs (DKO; Figures 1A and 1B), indicating that the autophagic response requires the presence of multidomain pro-apoptotic proteins. Surprisingly, BIM, tBID and PUMA induced more robust LC3 lipidation (Figure 1A) and accumulation of GFP-LC3-positive vesicles (Figure 1B) in *Bax-/-* MEFs compared to their *Bak-/-* counterparts, arguing that BAK plays a more predominant role in this experimental system. Similar results were obtained using the pro-apoptotic chemicals staurosporine (STS; Figures S1B and S1C) or mitoxantrone (MTX; Figures S1D and S1E), although the data were not as clean owing to high background activity in DKO and *Bak-/-* MEFs (see Figures S1B-E). This is likely due to stimulation of different stress pathways by the toxic drugs that activate a heterogeneous autophagic response including different components. Notably, neither DKO nor *Bak*-deficient MEFs showed alterations in basal (Figures S2A and S2B) or canonical (Figures S2C and S2D) autophagy, indicating that their inability to induce autophagy in this setting is not due to a defective autophagic machinery. In addition, the extent of cytochrome c release in response to BIM and tBID was comparable in *Bak-/-* and *Bax-/-* cells (Figure S3A), suggesting that their differential autophagic response is unrelated to mitochondrial outer membrane permeabilization (MOMP). Given their dissimilar autophagic response, we reasoned that *Bak-/-* and *Bax-/-* cells might provide a suitable system to study the functional consequences of autophagy induced during apoptotic cell death.

**Figure 1.**
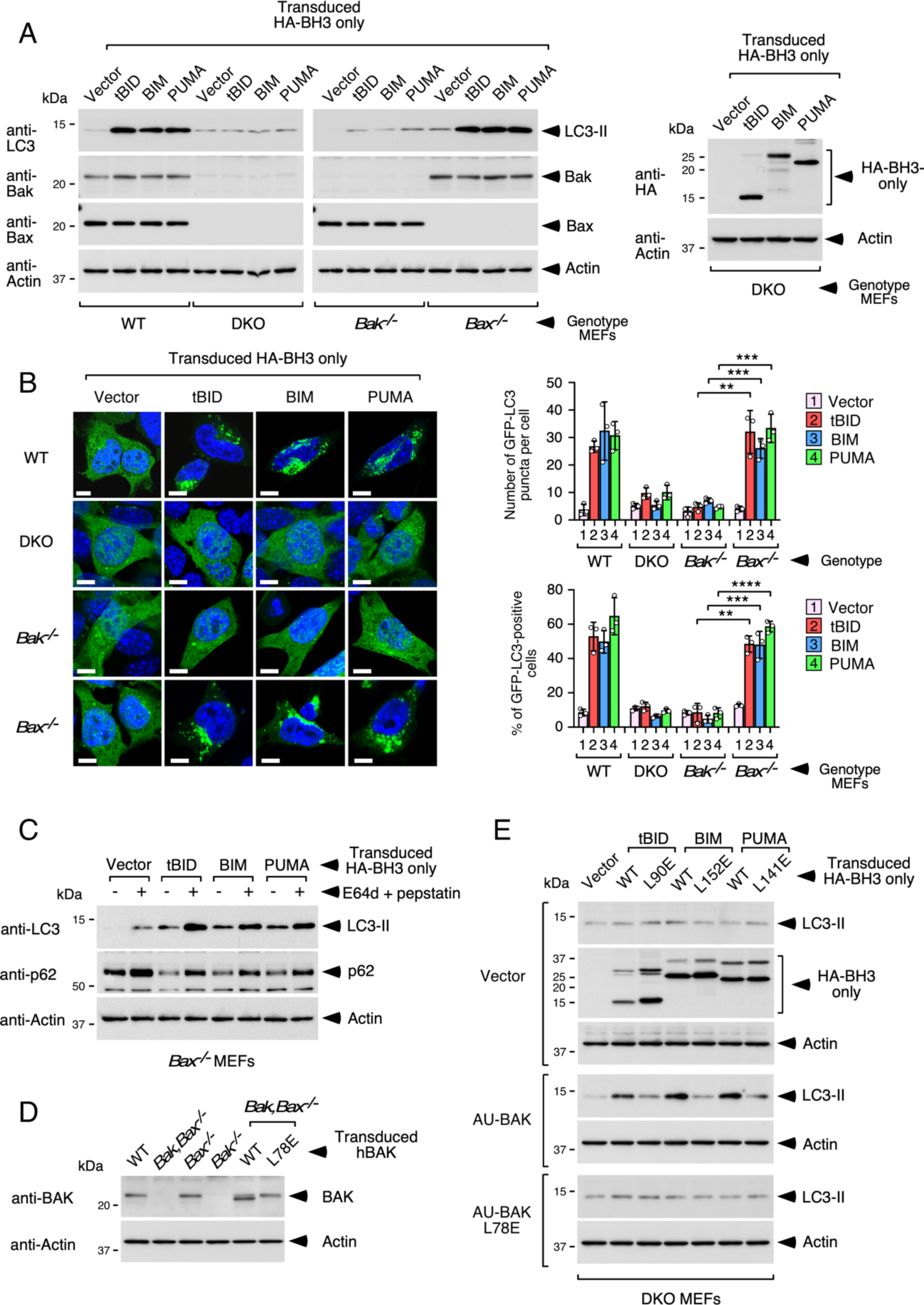
BH3-only activators induce BAK-dependent autophagy. **(A)** The indicated MEFs were transduced with retroviruses expressing the shown BH3-only molecules and treated with 25 μM zVAD.fmk 7 h later. Cells were lysed 22 h post-transduction for Western blot against the shown molecules. Expression levels of HA tagged tBID, BIM and PUMA were measured in DKO cells to minimize the impact of cell death (right panel). **(B)** MEFs expressing GFP-LC3 were retrovirally transduced as in **A** and fixed 22 h later for microscopy. Shown are representative confocal pictures (left panel), quantification of the number of GFP-LC3 puncta per cell (right panel, top) and the percentage of cells showing activated GFP-LC3 (right panel, bottom). Graphs represent mean values -/+ s.d. of triplicate experimental points (n = 3 microscopy fields including at least 25 cells per field; **P<0.01, \*\*\**P*<0.001, \*\*\*\**P*<0.0001 Student’s *t*-test). **(C)** *Bax-/-* MEFs were retrovirally transduced as in **A** and treated with E64d and pepstatin (10 μg/ml each) for the last 16 h of culture. Cells were lysed 22 h post-transduction for Western blot using the indicated antibodies. **(D)** BAK expression in reconstituted DKO MEFs. The indicated MEFs along with DKO MEFs retrovirally transduced with WT or L78E human BAK versions were lysed for Western blotting. **(E)** The reconstituted MEFs shown in **D** were retrovirally transduced as in **A** and lysed 22 h later for Western blot against the shown molecules.

### Unconventional features of the autophagic route activated by BAK

To study the nature of the autophagic response, we focused on *Bax-/-* MEFs. Previous publications show that absence of caspase activity during apoptosis allows activation of the STING signaling pathway(*42*), which in turn is able to activate autophagy(*43*)(*44*). However, expression of BIM and tBID caused substantial LC3 lipidation even without zVAD.fmk treatment (Figure S3B). In addition, while we observed strong STING phosphorylation caused by both BIM and tBID in *Bax-/-* cells (Figure S3C), STING depletion did not blunt LC3 lipidation induced by both effectors (Figure S3C). These results argue that the STING route is unlikely to mediate the autophagic response triggered during the course of apoptotic death in this experimental system.

LC3 lipidation caused by BIM, tBID and PUMA in *Bax-/-* cells reflects an enhanced autophagic flux, as the induced levels of LC3-II were further stimulated by lysosomal inhibition (Figure 1C) and accompanied by reduced p62 expression (Figure 1C). Reconstitution of DKO cells with AU-BAK (Figure 1D) restored the LC3 lipidation response (Figure 1E), indicating that BAK is the molecular entity that mediates autophagy in this system. Pathway activation requires the BH3 domains of both BAK and the BH3-only proteins, since mutant forms where a critical L in the BH3 motif was changed to E(*45*), were inactive (Figures 1D and 1E). Knock-down of the core autophagic effectors ATG7 or ATG16L1 via siRNAs inhibited LC3 activation by BIM, tBID and PUMA (Figure S4A). However, depletion of upstream initiators of canonical autophagy (VPS34, ATG14) had no effect (Figure S4B), suggesting that the pathway is atypical and therefore probably different from a previously described autophagic response triggered after OMM permeabilization that requires the conventional route mediated by AMPK and ULK1(*46*).

Published evidence shows that the WD40 domain (WDD) of ATG16L1 mediates unconventional autophagic activities where LC3 becomes lipidated in intracellular membranes unrelated to canonical autophagosomes(*33*)(*40*)(*41*)(*39*). *Bax-/-* MEFs depleted of ATG16L1 using CRISPR/Cas9 and reconstituted with ATG16L1 lacking the WDD (Nt; residues 1-299; Figures 2A and 2B) showed unaffected basal (Figure 2C, left panel) and canonical (Figure 2C, right panel) autophagy, but displayed reduced LC3 activation in response to BH3-only expression (Figures 2D and 2E) or MTX (Figures 2F and 2G). Similar results were obtained in *Atg16l1-/-* MEFs reconstituted with full-length ATG16L1 or just the N-terminal domain (Figures S4C-E), and also in *Atg16l1-/-* HCT116 colon carcinoma cells expressing with full-length ATG16L1 or separate N-and C-terminal domains (split ATG16L1; Figures S4F-H). We previously described that *Atg16l1-/-* MEFs expressing the Nt domain of ATG16L1(*38*) and HCT116 cells harboring split ATG16L1(*41*) cannot activate unconventional autophagy mediated by the WDD (in the latter case because the sensing (WDD) and effector (Nt region) domains are uncoupled), but are fully able to sustain basal and canonical autophagy(*38*)(*41*). Together, these results indicate that a large portion of the autophagic response triggered during apoptosis is unconventional, as it does not require some of the upstream effectors of canonical autophagy (VPS34, ATG14) and is mediated by the C-terminal WD40 domain of ATG16L1.

**Figure 2.**
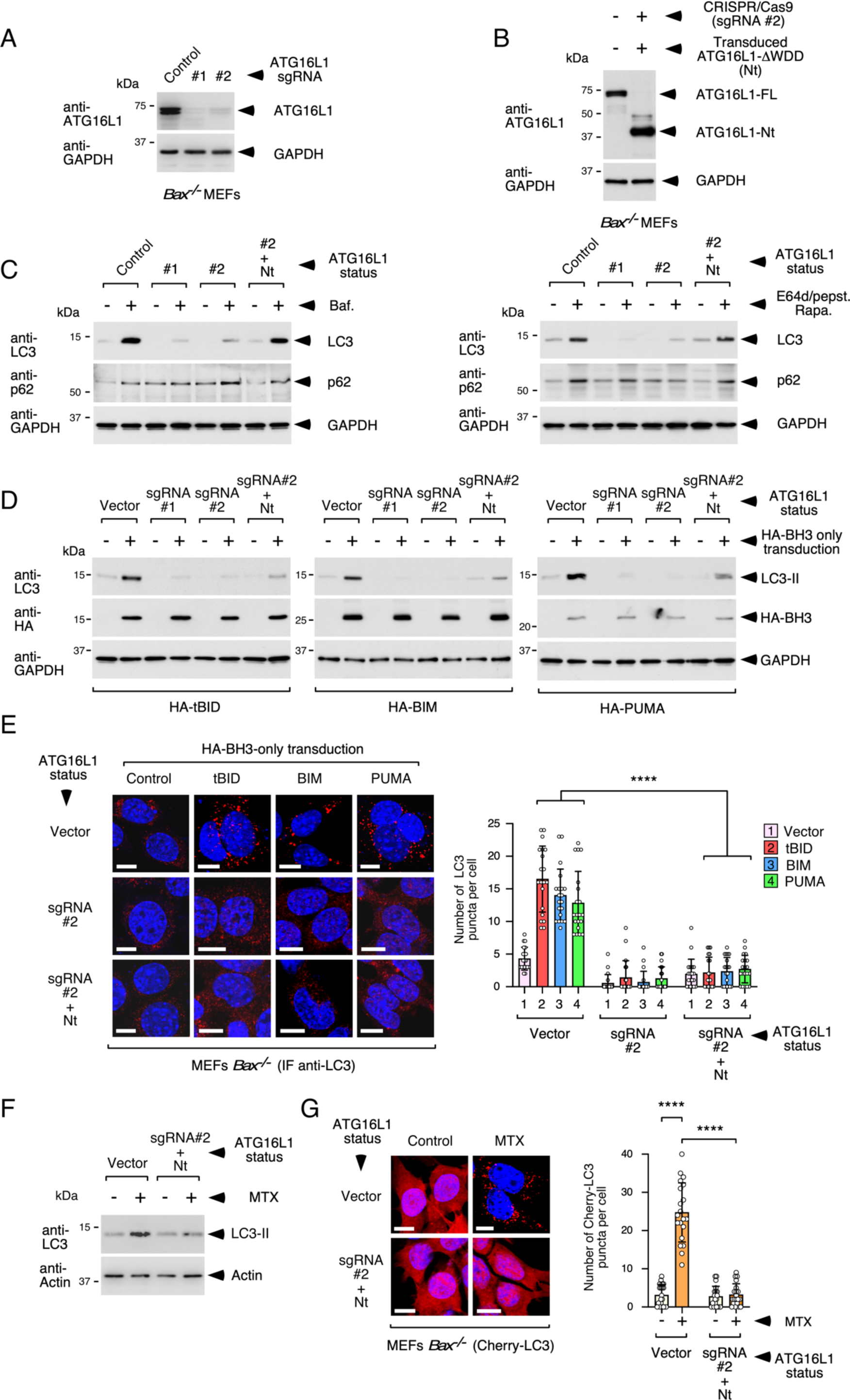
The autophagic response triggered in BAX-deficient cells requires the WD40 domain of ATG16L1. **(A)** *Bax-/-* MEFs were transduced with CRISPR/Cas9 lentivirus expressing the indicated sgRNAs against *Atg16l1* (#1 and #2), selected in puromycin and lysed for Western blot. **(B)** Cells targeted by sgRNA #2 were transduced with a retrovirus expressing the ATG16L1 Nt region (1-299), and lysed along CRISPR/Cas9 control cells for Western blot. **(C)** The engineered *Bax-/-* MEFs shown in **A** and **B** were treated with bafilomycin (80 nM; left panel) or rapamycin (2 μM) plus E64d/pepstatin (10 μg/ml each; right panel) for 8 h and lysed for Western blot. **(D)** The engineered *Bax-/-* MEFs shown in **A** and **B** were retrovirally transduced with the shown BH3-only activators, supplemented with 25 μM 7 h later, and lysed 20 h post-transduction for Western blot against the indicated molecules. **(E)** The indicated *Bax-/-* MEFs engineered for ATG16L1 expression were transduced as in **D** and fixed for anti-LC3 immunofluorescence (red) 17 h later. Shown are representative confocal pictures (left) and quantification of the number of LC3-positive puncta per cell (right). The graph shows mean values -/+ s.d. (n = 20 cells; \*\*\*\**P*<0.0001 Student’s *t*-test). **(F)** The indicated *Bax-/-* MEFs engineered for ATG16L1 expression were treated with 2 μM MTX in the presence of 25 μM zVAD.fmk and lysed 14 h later for Western blot against the shown molecules. **(G)** The indicated engineered *Bax-/-* MEFs stably expressing Cherry-LC3 were treated as in **F** and processed for microscopy 14 h later. Shown are representative confocal pictures (left) and quantification of the number of Cherry-LC3-positive puncta per cell (right). The graph shows mean values -/+ s.d. (n = 20 cells; \*\*\*\**P*<0.0001 Student’s *t*-test).

Confocal microscopy studies carried out in *Bax-/-* MEFs expressing GFP-LC3 and mitochondrial RFP (mRFP) showed that most of the LC3-positive vesicles generated by BH3-only expression or MTX do not contain mRFP (Figure S5A), arguing that the unleashed autophagic reaction does not reflect mitophagy. Mitophagic events were sporadically and distinctively observed in the same samples as a GFP-LC3-positive signal that grows tightly around, and encloses, the mitochondrial marker mRFP (Figure S5B; less than 1% of the GFP-LC3 positive puncta). In addition, while LC3 induction was evident 16 h post-transduction (Figure S5C), the levels of the mitochondrial marker VDAC1 remained unchanged in *Bax-/-* cells 24 h after BH3-only transduction (Figure S5C), indicating that there is no significant mitochondrial degradation in these conditions. Although most GFP-LC3-positive vesicles were isolated in the cytoplasm, some of the GFP-LC3-positive puncta localized in close apposition to the mitochondrial surface to the point of overlapping with mRFP to produce a yellow signal (insets in Figure S5A).

Closer inspection by electron microscopy revealed that the LC3-positive vesicles generated by BIM concentrate in a narrow plane of the perinuclear area (Figure 3A, left panels). For the most part, these vesicles presented a single membrane (Figure 3A, left panels, insets), so they are different from canonical double-membrane autophagosomes. Interestingly, some mitochondria were directly labelled with a GFP-LC3 signal that was not associated to other LC3-positive membranous structures (Figure 3A, right panel, top), arguing that these are not mitophagic events. This signal likely accounts for the close apposition events between GFP-LC3 and mRFP-labelled mitochondria detected in the confocal microscopy studies (see Figure S5A). Consistent with the confocal data, we detected rare mitophagic events that were easily identifiable (Figure 3A, right panel, middle). Intriguingly, tubular compartments consistent with endoplasmic reticulum (ER) cisternae were also sporadically labelled with GFP-LC3 (Figure 3A, right panel, bottom), suggesting that the approximately 15% of BAK that is targeted to reticular membranes in MEFs(*47*) may also be able to transmit autophagic signals.

**Figure 3.**
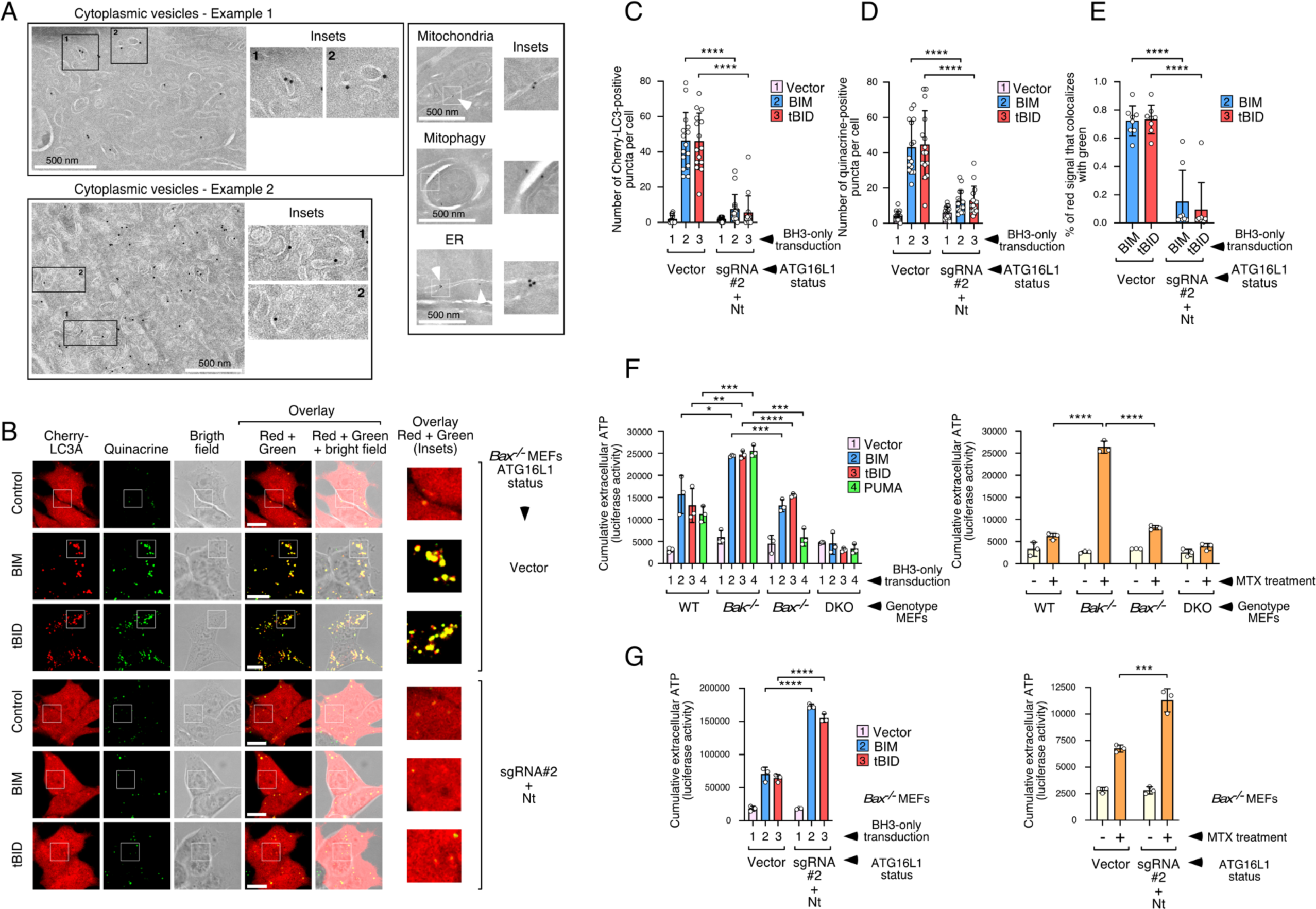
The unconventional autophagic response involves quinacrine-positive, single-membrane vesicles, and inhibits ATP release during cell death. **(A)** *Bax-/-* MEFs expressing GFP-LC3 were transduced with BIM, treated 7 h later with 25 μM zVAD.fmk, and processed 17 h post-transduction for anti-GFP immunoelectron microscopy. Shown are two examples of areas of accumulation of GFP-LC3-positive, single-membrane vesicles (left panels). Panels on the right display examples of mitochondria labelled with GFP-LC3 (top), mitochondria engulfed by GFP-LC3-positive membranes (mitophagy; middle) and GFP-LC3-positive tubules consistent with endoplasmic reticulum (ER) cisternae (bottom). **(B)** *Bax-/-* MEFs engineered for ATG16L1 expression and expressing Cherry-LC3 were transduced with BH3-only molecules and treated with 25 μM zVAD.fmk 7 h later. Cells were stained with quinacrine 17 h post-transduction and analyzed *in vivo* by confocal microscopy. Shown are representative confocal pictures. **(C-E)** Quantification of the phenotypes shown in **B**. Graphs display mean values -/+ s.d. of the number of Cherry-LC3-positive puncta per cell (n = 15 cells; **C)**, the number of quinacrine-positive puncta per cell (n = 15 cells**; D)** or the percentage of Cherry-LC3-positive signal colocalizing with quinacrine-positive signal (n = 8 cells; **E**). \*\*\*\**P*<0.0001 Student’s *t*-test. **(F)** The shown MEFs were transduced with BH3-only molecules for 22 h (left), or treated with MTX (2 μM) for 24 h (right), and the levels of extracellular ATP measured at 14 and 22 h (BH3-only molecules) or for 16 and 24 h (for MTX). Graphs show mean values -/+ s.d. of cumulative data resulting from the sum of the luciferase activity units obtained at both time points from triplicate experimental points (n = 3; \**P*<0.05, \*\**P*<0.01, \*\*\**P*<0.001, \*\*\*\**P*<0.0001 Student’s *t*-test). (G) *Bax-/-* MEFs engineered for ATG16L1 expression were transduced with BH3-only molecules (left) or treated with MTX (right), and the levels of extracellular ATP measured and displayed as in **F.**

### BAK-induced unconventional autophagy inhibits ATP release during apoptosis

*Bak-/-* and *Bax-/-* cells died to a similar extent in response to BH3-only activators (Figure S5D) or MTX (Figure S5E), and BIM, tBID and MTX induced comparable levels of cell death in cells expressing full-length ATG16L1 or just the Nt domain (Figures S5F-H). Therefore, the physiological role of the atypical autophagic response in this setting seems unrelated to the ability to modulate apoptosis that autophagy has in a variety of cell death paradigms(*23*). In search for a possible function, we found that most LC3-positive vesicles generated in WT and *Bax-/-* MEFs by BH3-only expression (Figures S6A-D) and MTX (Figures S7A-D) were stained by quinacrine, a lysosomotropic dye that emits green fluorescence in the presence of high ATP concentrations(*15*). The number of double LC3/quinacrine-positive and quinacrine-positive vesicles was reduced in treated *Bak-/-* MEFs (Figures S6A-D and S7A-D) and, notably, also in engineered *Bax-/-* cells lacking the WD40 domain of ATG16L1 (Figures 3B-E for BH3-only, and Figures S7E-H for MTX), raising the notion that the LC3-positive vesicles generated by the atypical BAK-dependent autophagic pathway could be sequestering ATP. Previous reports showed that autophagy activated during immunogenic apoptosis promotes the accumulation of ATP inside canonical autophagosomes to be delivered for secretion(*21*). Unexpectedly, however, we found that *Bak-/-* cells (where the autophagic pathway is impaired) secrete more ATP during apoptosis compared to WT and *Bax-/-* cells (where the autophagic pathway is fully active), both in response to BH3-only expression (Figure 3F, left) and MTX (Figure 3F, right), and retain less ATP remaining in the apoptotic bodies than *Bax-/-* cells (Figures S8A and S8B). These data argue that the atypical autophagic response induced by BAK may in fact prevent ATP secretion. Consistently, engineered *Bax-/-* cells lacking the WD40 domain of ATG16L1 showed increased ATP liberation in response to BH3-only (Figure 3G, left) and MTX (Figure 3G, right), and reduced ATP levels remaining in the apoptotic bodies (Figures S8C and S8D), compared to their control counterparts. Similar observations were made in *Atg16l1-/-* MEFs reconstituted with full-length ATG16L1 or just the Nt domain (Figures S8E and S8F for BH3-only molecules, and S8G and S8H for MTX), and also in *ATG16l1-/-* HCT116 cells harboring full-length ATG16L1 or separated Nt and Ct domains; Figure S8I for BH3-only molecules, and S8J for MTX). Therefore, data obtained in different experimental systems reveal the existence of an unconventional autophagic route that emanates from activated BAK and prevents ATP secretion during apoptosis, likely by sequestering it into LC3-positive vesicles.

### The latch domain of BAK induces LC3 lipidation

In search for the molecular mechanism that links activated BAK to the induction of unconventional autophagy, we found that a mutated version of BAK lacking the BH3 proapoptotic domain (BAK-ΔBH3) is able to induce LC3 lipidation in a straight overexpression assay (Figures S9A and S9B) without significantly inducing cell death compared to wild-type BAK (Figure S9C). In contrast, BAX-ΔBH3 was unable to activate LC3 (see Figures S9A and S9B). These results suggest that BAK has an intrinsic autophagic activity that might reflect its ability to induce unconventional autophagy in response to BH3-only effectors. We reasoned that characterization of this function could help identify downstream molecular machinery engaged by activated BAK to induce the noncanonical pathway.

In the native structure of BAK, the BH3 domain is deeply buried into a hydrophobic core(*48*), so we reasoned that BAK-ΔBH3 could be partially unfolded. Since this mutant strongly induces autophagy, there is a chance that its tertiary structure is not required for the autophagic function, and therefore it might be possible to identify a region of the primary sequence that retains the activity. To this end, we initially broke down the BAK open reading frame into portions that make structural sense. Upon activation, BAK sequentially unfolds, first, to expose its N-terminal region (helix 1) and, second, to separate the core (helices 2-5) and latch (helices 6-8) domains before dimerizing/multimerizing to generate OMM pores(*49*)(*50*)(*4*). Since BAK needs the hydrophobic C-terminal region (helix 9) for mitochondrial targeting, we generated constructs expressing these different domains fused to the TM helix for mitochondrial localization (Figure 4A). Overexpression studies indicated that the autophagic activity is exclusively retained by the latch domain (G146-G186; Figures 4B and 4C, and scheme in Figure 4D). Expression of this region did not cause cell death (Figure S9D), indicating again that the autophagic route can be segregated from the induction of apoptosis. *Atg16l1-/-* HCT116 cells reconstituted with separated N- and C-terminal domains of ATG16L1 did not respond to latch expression with LC3 lipidation (Figure S9E), thus confirming the unconventional nature of the autophagic pathway. The latch domain was able to induce LC3 lipidation in HCT116 cells lacking both BAK and BAX (Figure S9F, left and right panels), indicating that its autophagic activity is not channeled through activation of these endogenous effectors.

**Figure 4.**
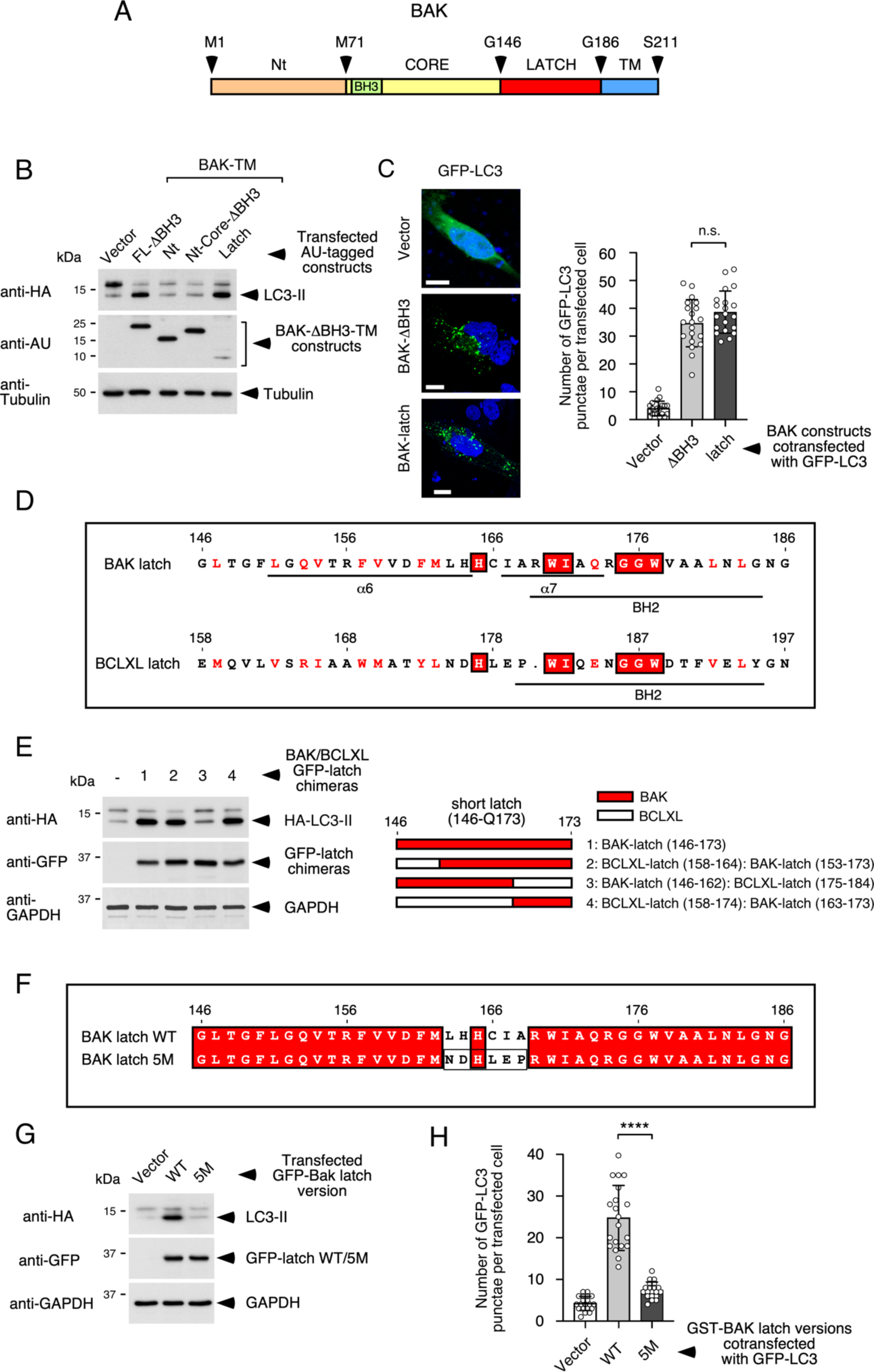
The BAK latch domain suffices to induce LC3 activation. **(A)** Scheme of human BAK showing relevant amino acid positions used for the structure-function studies in this figure. **(B)** HEK-293T cells were transfected with plasmids expressing the shown AU-tagged BAK-ΔBH3 constructs (Nt: M1-M71; Nt-core-ΔBH3: M1-G146 lacking the BH3 region (G72-R88); latch: G146-G186; all including the TM domain: P187-S211), and HA-LC3. Cells were lysed 36 h later and processed for Western blot against the indicated molecules. **(C)** Hela cells were transfected with the indicated AU-tagged BAK constructs and GFP-LC3. Cells were processed for microscopy 36 h later. Shown are representative confocal pictures (left) and quantification of the number of GFP-LC3-positive puncta per cell (right). The graph shows mean values -/+ s.d. (n = 20 cells; n.s. *P*>0.05 Student’s *t*-test). **(D)** Aligned sequences of the BAK and BCLXL latch domains showing functionally conserved (red font) and identical (red boxes) amino acids. The positions of the BH2 domains and the α6 and α7 helices of BAK are indicated. **(E)** HEK-293T cells were transfected with the indicated GFP-tagged BAK/BCLXL latch chimeric constructs (right panel), along with a plasmid that expresses HA-LC3. Cells were lysed 36 h later and processed for Western blot (left panel). **(F)** Scheme showing wild-type and 5M BAK latch domain versions. Red boxes include identical residues and boxed amino acids in the 5M version indicate the residues of BCLXL (ND-LEP) that substitute those present in BAK (LH-CIA). **(G)** HEK-293T cells were transfected with plasmids expressing WT and 5M GFP-tagged BAK latch constructs (as indicated) along with HA-LC3. Cells were lysed 36 h later and processed for Western blot. **(H)** Hela cells were transfected with the indicated plasmids expressing WT and 5M GST BAK latch versions and a plasmid that expresses GFP-LC3. Cells were fixed 36 h later, processed for microscopy and quantified for the number of GFP-LC3-positive puncta per cell. The graph shows mean values -/+ s.d. (n = 20 cells; **** *P*<0.0001 Student’s *t*-test).

Systematic N- and C-terminal deletion studies showed that the minimal latch version that retains the autophagic activity lies between amino acids G146-Q173 (α6-α7) (Figures S9G, top and bottom panels, and scheme in Figure S9H). An equivalent region of the multidomain anti-apoptotic homologue BCLXL (E158-E184; Figure S9H) was inactive even when fused to the TM domain of BAK (Figure S9I, left panel), as it was the full-length BCLXL latch domain (E158-N197; Figure S9I, right panel). Studies using chimeric constructs between the minimal BAK and BCLXL latch fragments pointed to a region of the former that is critical for the activity (residues L163-Q173; Figure 4E, left and right panels). This region overlaps with the BH2 domain of BAK that is highly homologous between BCL2 family members, so we hypothesized that the most relevant residues might be those excluded from the BH2 sequence (L163-A168; see scheme in Figure 4D). Consistently, changing these residues to those present in BCLXL (BAK-latch-5M; Figure 4F) impaired the autophagic activity of both the BAK latch domain (Figure 4G and 4H) and BAK-ΔBH3 (Figures S9J and S9K). Attempts to express a 5M version of BAK in DKO MEFs were unsuccessful due to instability and reduced expression levels of the mutant protein (not shown).

### A complex including PHBs and STOM links the latch and WD40 domains

To identify proximal signaling machinery that might be engaged by the latch domain to induce unconventional autophagy, fusion constructs between GST and wild-type or mutant (5M) latch versions were transfected into HEK-293T cells and purified to identify co-precipitating proteins by mass spectrometry. Out of a total of 661 proteins identified (Source Data 1), 239 were preferentially present in the wild-type sample at least by one more identified peptide compared to the 5M version (Source Data 2). We further selected a short list of 35 candidates that fulfilled the arbitrary threshold of being overrepresented by more than 4 identified peptides in the wild-type sample (highlighted candidates in Source Data 2 file). Members of the importin/exportin family of transporters across the nuclear membrane were prominently present in this short list (Figure S10A), but co-immunoprecipitation studies between GST-latch and the highest-ranked family candidate (EXPORTIN-4) revealed no signs of interaction (Figure S10B). Intriguingly, the short list of candidates also included 4 members of the prohibitin and stomatin family: prohibitins 1 and 2 (PHB1, PHB2), stomatin (STOM) and stomatin-like protein 2 (SLP2)(*51*), as well as the mitochondrial porin VDAC1 (see Figure S10A and Figure 5A). PHB1 and PHB2 are multifunctional, heterodimeric proteins located in the mitochondrial intermembrane space and critical for mitochondria metabolism and function(*52*). STOM is a membrane-associated protein located in lipid rafts that regulates channel activity(*51*), and SLP2 interacts with PHBs and controls the stability of respiratory complexes and mitochondrial fusion(*51*). VDAC1 was also selected because the family member VDAC2 was previously described as a BAK interactor(*53*). Co-immunoprecipitation studies confirmed the specific interaction between the latch domain and PHB1/2, STOM and VDAC1, both in a co-transfection modality (Figure S10C, top, middle and bottom panels, respectively) and also after expression of GST-latch and detection of the coprecipitating endogenous proteins (Figure S10D, top, middle and bottom). However, SLP2 did not bind the latch domain in any of the experimental configurations (cotransfected, Figure S11A or endogenous, Figure S11B). CRISPR/Cas9-mediated depletion of PHB2 or STOM (Figures S11C and S11D), but not that of VDAC1 or SLP2 (Figures S11E and S11F), inhibited LC3 activation caused by expression of the latch domain, demonstrating that they mediate the autophagic response in this context. In light of these results, VDAC1 and SLP2 were excluded from further studies. Both PHB1/2 and STOM specifically co-precipitated with the latch domain after co-expression (Figure S12A, left and right panels), arguing that the three contenders might be able to assemble into a complex nucleated by the latter. To establish the hierarchy of such possible complex we performed co-precipitation studies between GST-latch and endogenous PHBs and STOM in cells depleted for the different candidates using the CRISPR/Cas9 system. Depletion of STOM did not alter the ability of the latch domain to bind endogenous PHBs (Figure S12B), but reduction of PHB levels impaired co-precipitation between GST-latch and endogenous STOM (Figure S12C). These results indicate that the interaction between STOM and the latch domain is probably bridged by PHBs. Consistently, PHB1/2 were found in STOM-GST precipitates (Figure S12D), arguing that these proteins can bind each other. The interaction between PHBs and the latch region is probably direct, since a pre-purified wild-type version of the latch domain fused to GST was able to pull-down both PHBs from a crude cell lysate, whereas the 5M mutant was inactive (Figure S12E). This result also indicates that the failure of the 5M mutant to bind PHBs is not secondary to a possible inability to traverse the OMM to reach both PHBs.

**Figure 5.**
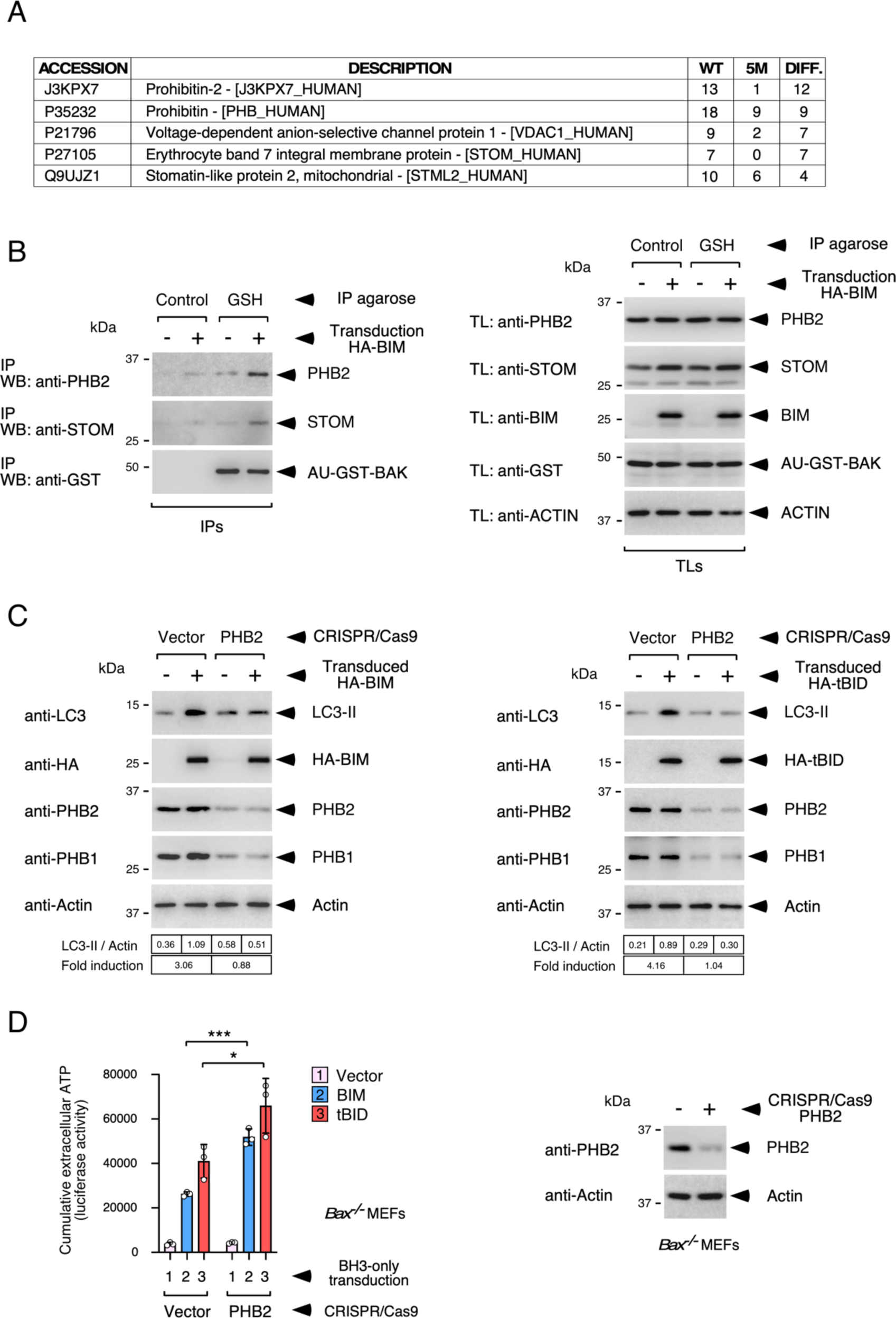
A BAK-nucleated complex including PHB1/2 and STOM mediates BAK-induced autophagy and inhibits ATP secretion during apoptosis. **(A)** Selected list of prohibitin-family proteins and VDAC1 identified as potential latch interactors. The number of peptides identified in the WT and 5M proteomics samples, and their difference between both samples (DIFF.), are indicated. **(B)** Co-immunoprecipitation assay showing interaction between activated GST-BAK and PHB2/STOM in response to BIM. DKO MEFs depleted of CASP3 and CASP9 and expressing GST-BAK (and also human PHBs and STOM for increased detectability) were retrovirally transduced with HA-BIM, and then crosslinked with 1% paraformaldehyde and lysed 15 h later. Lysates were incubated with control or GSH-agarose beads and the resulting precipitates subjected to Western blot (left panels, IPs). Expression levels of all contenders in total protein lysates are shown on the right panel (TLs). **(C)** PHBs mediate the autophagic response induced by BIM (left panel) and tBID (right panel). *Bax-/-* MEFs were transduced with the shown CRISPR/Cas9 constructs and, 40 h later, with the indicated BH3-only molecules. Cells were treated with 25 μM zVAD.fmk 7 h later and lysed 20 h post-transduction for Western blot. Densitometric quantifications are shown at the bottom. **(D)** PHBs inhibit ATP secretion during BH3-only-induced apoptosis. *Bax-/-* MEFs were transduced with the shown CRISPR/Cas9 constructs and the indicated BH3-only molecules (as in **C**) for 22 h, and the levels of extracellular ATP were measured at 14 and 22 h. Graph shows mean values -/+ s.d. of cumulative data resulting from the sum of the luciferase activity units obtained at both time points from triplicate experimental points (n = 3; \**P*<0.05, \*\*\**P*<0.001, Student’s *t*-test). Control points were lysed 48 h post-CRISPR/Cas9 transduction for Western blot to determine PHB2 depletion (right panel).

Co-immunoprecipitation assays carried out in DKO MEFs reconstituted with GST-BAK (Figure 5B, left and right panels) or untagged BAK (Figure S13A, left and right panels) (and also expressing human PHBs and STOM for increased detectability), showed that BAK activated by BIM is able to bind PHB2 and STOM. Considered together, these results show that, upon activation by BH3-only effectors, BAK unfolds to expose the latch domain for binding to PHBs/STOM.

CRISPR/Cas9-mediated depletion of PHB1/2 inhibited LC3 activation by BIM and tBID in *Bax-/-* MEFs (Figure 5C), and caused increased ATP secretion in both BAX-deficient (Figure 5D) and wild-type MEFs (Figure S13B) in response to the same activators, thus confirming that the unconventional autophagic response triggered during apoptosis to inhibit ATP release is mediated by PHBs. Therefore, as observed in the overexpression experiments carried out with the isolated latch domain (see Figures S10C-D and S11C), BAK activated by BH3-only inducers also physically and functionally interacts with PHBs to induce unconventional autophagy. These results argue that the functional properties of the isolated latch domain faithfully recapitulate the activity of the native protein in this pathway.

We previously published a proteomics study to identify proteins that may interact with the WD40 domain of ATG16L1(*54*). Notably, both PHB1 and STOM were included in the candidate list (Figure S14A and Source Data 3), suggesting that both proteins could be WDD interactors. Consistently, PHB1/2 (Figure S14B) and STOM (Figure S14C) were able to bind a GST-ATG16L1-320-607 (WDD) construct in co-immunoprecipitation studies. Since the autophagic activity of the latch domain requires a functional WDD (see Figure S9E) we hypothesized that the PHBs/STOM complex could help link the latch domain to ATG16L1 through the WDD. Consistent with this view, while the latch region is by itself unable to interact with ATG16L1 (Figure S14D, left and right panels), the presence of PHBs and STOM facilitated the presence of a GFP-latch fusion protein in GST-ATG16L1-(320-607) (Figure S14E, left and right panels) precipitates, arguing that the PHB/STOM complex engaged by the latch domain may help recruit ATG16L1 to mitochondrial membrane sites. In agreement with this notion, GFP-ATG16L1 is recruited to the mitochondrial surface in *Bax-/-* MEFs transduced with BIM or tBID (Figure S15A and S15B). Together, these data argue that BAK, once activated by BH3-only molecules, assembles a complex including PHBs and STOM that interacts with ATG16L1 through the WDD to facilitate the induction of unconventional autophagy and inhibition of ATP release.

### The unconventional autophagic response inhibits IL-1β secretion by BMDMs

Previous evidence shows that ATP released during cell death promotes activation and differentiation of the phagocytes that remove the cellular remains(*55*). These results suggest that the unconventional autophagic route stimulated by BAK might inhibit phagocyte activation by suppressing ATP mobilization. Consistently, we found that murine bone marrow-derived macrophages (BMDMs) pre-stimulated with LPS secrete increased amounts of IL-1β when they are co-incubated with engineered BAX-deficient apoptotic cells lacking the WDD (Figures 6A and 6B). Similar observations were made using reconstituted *Atg16l1-/-* MEFs expressing full-length ATG16L1 or just the Nt domain (Figure S15C), and also in *ATG16l1-/-* HCT116 cells harboring full-length or split ATG16L1 (Figure S15D), although the effect in the latter case was more discrete, perhaps because the stimulatory effect of the strain expressing FL ATG16L1 was particularly strong so that the contribution of increased ATP secretion may be less relevant. Treatment with two different inhibitors of the purinergic P2X7 receptors (KN-62 and JNJ-47965567) blunted IL-1β liberation caused by *Bax-/-* cells expressing ATG16L1-ΔWDD (Figure 6C), thus confirming that ATP is the molecular mediator that causes IL-1β secretion by BMDMs in this experimental context. These results indicate that the unconventional autophagic activity that emanates from BAK truly represses the immunogenic potential of cell death, and underscore the notion that the secreted ATP fraction, and not the pool that remains sequestered inside the cellular remains, is the one that has immunogenic properties.

**Figure 6.**
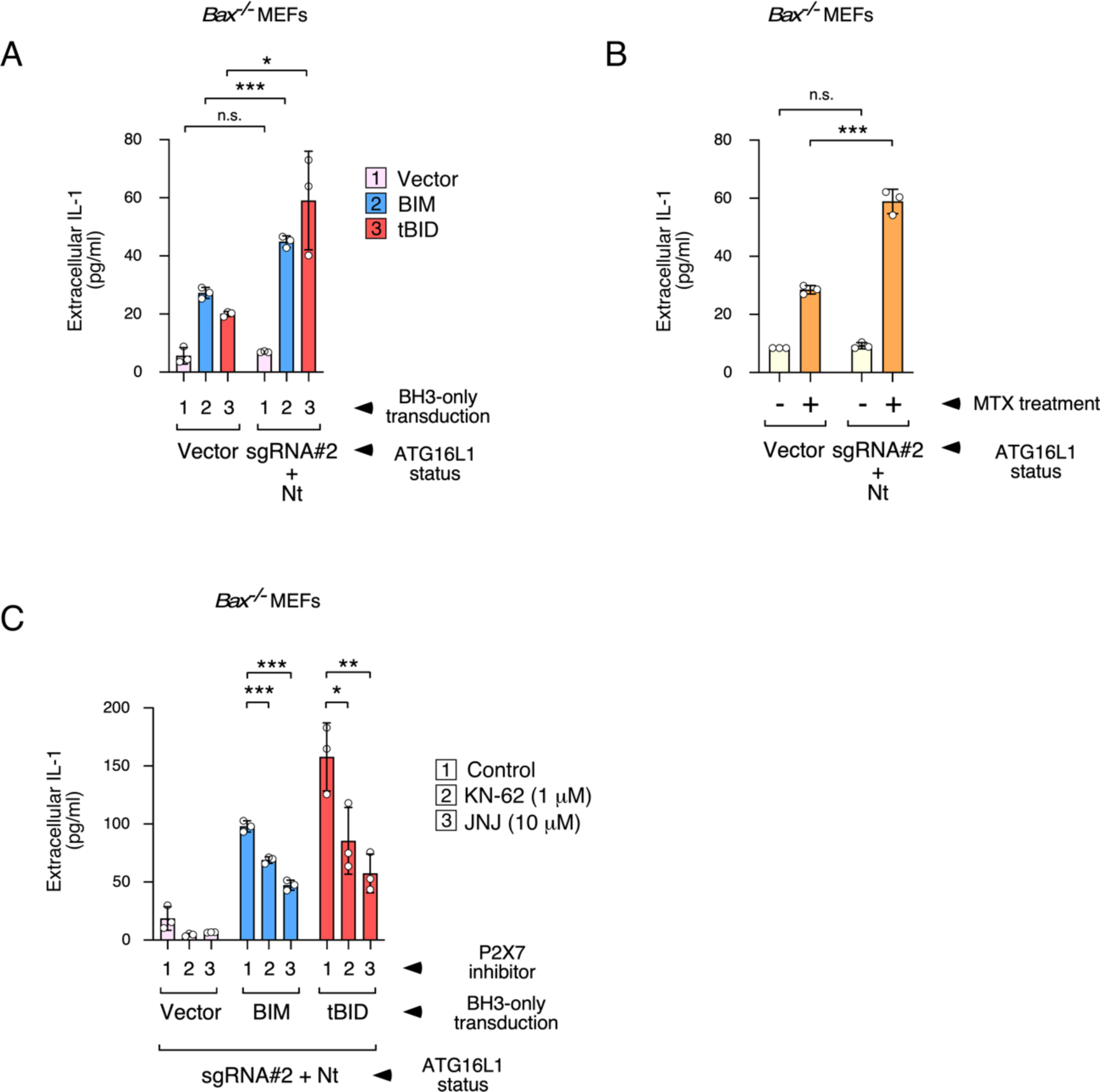
The unconventional autophagic response of apoptotic *Bax-/-* MEFs suppresses IL-1β release by co-cultured LPS-activated BMDMs. **(A)** *Bax-/-* MEFs engineered for ATG16L1 expression (as indicated) were transduced with retroviral constructs expressing the shown BH3-only molecules and, 7 h later, washed and supplemented with 3 μM of the ecto-ATPase inhibitor ARL67156. Apoptotic bodies were collected 20 h post-transduction and added to differentiated BMDMs pre-activated for 4 h with 100 ng/ml of LPS. After 6 h, the culture medium was tested for IL-1β concentration by ELISA. Graphs show mean values of IL-1β concentration -/+ s.d. of triplicate experimental points (n = 3; n.s. *P*>0.05, \**P*<0.05, \*\*\**P*<0.001 Student’s *t*-test). **(B)** *Bax-/-* MEFs engineered for ATG16L1 expression (as indicated) were treated with 2 μM MTX and, 7 h later, washed and supplemented with 3 μM of the ecto-ATPase inhibitor ARL67156. Apoptotic bodies were collected 22 h after MTX treatment and added to differentiated BMDMs as in **A**. Graphs show IL-1β concentration data as in **A** (n = 3; n.s. *P*>0.05, \*\*\**P*<0.001 Student’s *t*-test). **(C)** *Bax-/-* MEFs engineered for ATG16L1 expression to harbor the ATG16L1 Nt domain (residues 1-299) were transduced with the indicated BH3-only molecules and washed 7 h later. The resulting apoptotic bodies were collected 20 h post-transduction and added to BMDMs pre-incubated for 45 min with the shown doses of the P2X7 channel inhibitors KN-62 and JNJ. After 6 h incubation in the presence of the same doses of the inhibitors, the culture medium was tested for IL-1β concentration. Graphs show IL-1β concentration data as in **A** (n = 3; * *P*<0.05, \*\**P*<0.01, \*\*\**P*<0.001 Student’s *t*-test).

## DISCUSSION

Our results reveal a novel signaling route that links the intrinsic apoptotic machinery to the induction of unconventional autophagy. This pathway is initiated by the multidomain pro-apoptotic effector BAK once it is activated by a defined subset of BH3-only molecules. The autophagic activity appears unrelated to the induction of cell death, since *Bak-/-* and *Bax-/-* MEFs release cytochrome c and die to a similar extent in response to the BH3-only inducers while showing dissimilar autophagic responses. Notably, BAK induces this route by binding the multifunctional mitochondrial effectors PHB1 and PHB2, and STOM, revealing the existence of a previously unappreciated interactome assembled by unfolded BAK to trigger alternative activities not directly linked to OMM pore formation. Since PHBs are located at the inner mitochondrial membrane facing the intermembrane space(*52*), our results may have implications regarding the topology of activated, unfolded BAK at the surface of the OMM. One view suggests that the latch domain remains flat on the OMM, whereas an alternative theory argues that it penetrates into the OMM to favor pore formation(*4*)(*6*)(*56*)(*57*). Since the latch domain likely needs to cross the OMM to bind both PHBs, our data seem to favor the second model. It is also conceivable that PHBs may change their location during apoptosis to become more accessible to the latch domain and STOM. While STOM has been described as a major lipid raft component(*58*), it has also been found in mitochondria-associated structures during viral infection(*59*), so a mitochondrial localization of a STOM fraction for binding to the BAK latch/PHB complex is plausible.

Our data sustain the idea that the molecular assembly formed by BAK, PHBs and STOM interacts with the WDD of ATG16L1 to facilitate formation of single-membrane unconventional autophagic vesicles. However, the exact origin of these vesicles (for instance, whether they are formed *de novo* or by fragmentation of an existing organelle) is unclear. Mitochondria have been shown to act as membrane donors for the generation of autophagic vesicles(*60*)(*61*), but we did not detect vesicular structures emanating from the mitochondrial surface in our electron microscopy studies. Mitochondria are also known to undergo a fragmentation process during apoptosis(*62*), but most of the GFP-LC3-positive vesicles detected in our assays were devoid of the mitochondrial marker mRFP, thus arguing against a direct mitochondrial origin. Interestingly, PHB2 is a receptor for selective autophagy of the inner mitochondrial membrane once the OMM has been damaged(*63*). This phenomenon may explain the direct labelling of apoptotic mitochondria with GFP-LC3 that we detected in both confocal and immunoelectron microscopy studies. However, these incipient events did not progress to mitophagy to a significant extent in our experimental system, so their relevance in the pathway that we describe here is unclear. While our results suggest that direct recruitment of ATG16L1 to the BAK/PHB/STOM complex helps generate the LC3-positive vesicles, it is possible that a diffusible element released by porated mitochondrial might also play a relevant role.

Previous publications show that MOMP, and the consequent decreased levels of ATP, activate autophagy through a canonical AMPK and ULK1-dependent pathway that degrades mitochondria(*46*). However, our data indicate that BAK- and WDD-induced autophagy is probably different from this activity, since it does not require upstream inducers of the canonical route (ATG14, VPS34), and it does not result in substantial mitophagy. In addition, previous results show that MOMP causes the release of mitochondrial DNA during apoptosis(*42*)(*64*)(*65*), which in turn can activate the STING pathway provided that caspase activity is blunted(*42*). Separate studies indicate that STING stimulation induces autophagy, although the nature of this response is unclear, with canonical(*43*) and unconventional components suggested(*44*). However, our results showing that BAK-induced autophagy occurs without the need of caspase inhibition, and also that STING depletion does not inhibit LC3 lipidation, appear to exclude a relevant role of STING in the pathway that we describe here.

We found that the novel autophagic route activated by BAK inhibits ATP release during apoptosis, thus repressing immunogenic cell death. Mechanistically, our results favor a model where ATP is sequestered inside single-membrane LC3-positive vesicles to prevent its secretion during the course of apoptosis. How ATP may become trapped into these vesicles is currently unknown, but other processes where ATP is sequestered into cytoplasmic vesicles have been described. For instance, canonical autophagosomes appear to accumulate ATP for secretion during immunogenic apoptosis induced by chemotherapeutic drugs(*18*)(*21*). Other autophagic activities have been shown to occur during apoptotic death to, for example, sustain ATP levels for successful execution of cell death(*66*), induce secretion of HMGB1(*17*) or promote mitophagy(*67*)(*46*)(*68*). While mitophagy could potentially contribute to inhibit ATP secretion by sequestering and degrading mitochondria, thus limiting the pool of ATP available for liberation, the low levels of mitophagy occurring in our experimental setting do not seem to support this idea. Therefore, the unconventional autophagic process that we describe here may be induced along with other autophagic responses during cell death, supporting a scenario where phenotypically-alike but functionally-different LC3-positive vesicles originated in various subcellular locations, containing different cargoes and performing different functions probably coexist. How these different mechanisms may overlap and interact with each other is an interesting question that remains to be investigated.

More generally, our results point to the idea that cell death immunogenicity is the default outcome of the apoptotic program that is actively repressed by the same machinery that promotes the cell’s suicide in order to favor an immunosilent process. This view implies that inhibition of unconventional autophagy may boost the immunogenic potential of cell death, an attractive property that could be used with therapeutic purposes as an anti-cancer strategy. In fact, since the autophagic route that regulates this activity has a number of atypical features, it could conceivably be inhibited without altering canonical autophagy, thus avoiding any unwanted side effects that may be associated to impairment of this critical housekeeping process.

## METHODS

### Cell lines and reagents

HEK293T and HeLa cells were purchased from the American Type Culture Collection. WT, *Bax-/-*, *Bak-/-* and DKO (*Bax-/-*, *Bak-/-*) SV40-transformed MEFs were obtained from Dr. Stanley Korsmeyer’s laboratory(*69*). *Atg16l1-/-* MEFs were kindly shared by Dr. Ramnik Xavier (Broad Institute MIT and Harvard, USA)(*70*), and were reconstituted via retroviral transduction with full-length HA-ATG16L1 or an AU-Nt fragment (1-299) fragment, as described(*38*). *ATG16L1-/-* HCT116 cells were obtained from Dr. David Boone’s group (Indiana University School of Medicine, USA)(*71*), and reconstituted with either full-length HA-ATG16L1 or AU-Nt (1-299) plus HA-Ct (300-607) fragments simultaneously, as previously described(*41*). Cells were cultured at 37° C and a humidified 5% CO_2_ atmosphere, in DMEM (Invitrogen, 52100-047) containing 10% heat-inactivated FBS (Invitrogen, 10437028) and 100 U/ml of penicillin/streptomycin (Invitrogen, 15140130). Chemicals: zVAD.fmk (Becton Dickinson, 550377), DAPI (Sigma, 10236276001), ARL67156 (Sigma, A265), etoposide (Sigma, E1383), E64D (EST, Calbiochem, 330005), pepstatin (Sigma, P4265), polybrene (Sigma, H9268), bafilomycin A1 (Sigma, B1793), rapamycin (Selleckchem, S1039), staurosporine (Sigma, S4400), mitoxantrone (MTX, Sigma, M6545), quinacrine (Sigma, Q3251), KN-62 and JNJ-47965567 (Tocris, 1277 and 5299, respectively).

### DNA constructs, transfections and retroviral transductions

Constructs were expressed from the mammalian expression vector Peak12 or its retroviral derivative Peak12-MMP. To express BAK in DKO MEFs, human BAK was cloned into a version of the retroviral Peak12-MMP vector containing an IRES-hygromycin cassette for selection in 200 μg/ml hygromycin B (Invitrogen, 10687010). The cDNAs for all BH3-only members, BAK, BAX and VDAC1 were amplified by PCR from a human T cell library and fully verified by sequencing. Human PHB1 and PHB2 cDNAs were purchased from Genescript (refs. OHu17487 and OHu16881, respectively), and could only be expressed to a significant level if co-transfected. Other cDNAs were kindly shared with us by other investigators: human PUMA (Dr. Bert Vogelstein, Johns Hopkins University, MD, USA), BCLXL (Dr. Craig Thompson, Memorial Sloan Kettering Cancer Center, NY, USA), STOMATIN (STOM, Mario Mairhofer, Univ of Applied Sciences, Austria), Flag-EXPORTIN-4 from Dirk Goerlich (Max Plack Institute, Gottingen, Germany) and Stomatin-like protein 2 (SLP2 from Dr. Joaquin Madrenas, UCLA, Los Angeles, USA). Tagged versions of all constructs were built by PCR. To generate BAK-ΔBH3 (G72-R88) and BAX-ΔBH3 (L59-N73) the BH3 domains were substituted by an EcoR1 site (EF frame) by PCR. BAK-ΔBH3 deletions (Nt: M1-M71; Nt-core-ΔBH3: M1-G146 lacking the BH3 region (G72-R88) and latch: G146-G186), were all fused at the N-terminus of the BAK C-terminal TM domain (P187-S211) and were generated by PCR. BAK-latch Nt and Ct deletions, BAK-BCLXL-latch chimeric constructs #2 (BCLXL(158-164)-BAK(153-173)), #3 (BAK(146-162)-BCLXL(175-184)), #4 (BCLXL(158-174)-BAK(163-173)), and ATG16L1 constructs (N-terminal domain, 1-299; C-terminal domains, 300-607, 231-607 and 320-607) were also generated by PCR. PCR oligonucleotides are shown in Table S1. All constructs were fully verified by sequencing. The BH3-domain mutants of BAK, BIM, tBID, PUMA, and the BAK latch 5M mutant was generated by site-directed mutagenesis using the mutagenic oligonucleotides shown in Table S1. Plasmids were expressing the baculovirus p35 apoptotic inhibitor, CASPASE-9 dominant-negative (C287A), mitochondrial RFP and GFP(*72*), and tagged LC3A constructs(*33*), were previously described.

Transfections were carried out using the jetPEI reagent (Polyplus, PPLU10-40) according to the instructions provided by the manufacturer. Retroviral transductions were done using virus-containing supernatants previously generated by co-transfecting HEK-293T cells with the relevant P12-MMP constructs and helper plasmids expressing gag-pol (pMD.gag-pol) and env (VSV-G; pMD-G). Supernatants were filtered against 0.45 μM, diluted with fresh medium (1:1) and supplemented with polybrene (8 μg/ml). Infections were done by spinning the resulting mix onto the target cells for 1 h at 2000 rpm, 32°C. In those experiments where cell death induced by BH3-only molecules had to be minimized, zVAD.fmk was added 7 h post-transduction.

### CRISPR/Cas9 and siRNAs

For protein depletion using the CRISPR/Cas9 system, we used the lentiCRISPRv2 vector (that includes a puromycin-selection cassette; a gift from Dr. Feng Zhang, Addgene, Ref. 52961), and its blasticidin or hygromycin derivatives (a gift from Dr. Brett Stringer, Addgene, Refs. 98293 and 98291, respectively). Guides for human and/or mouse PHB1, PHB2, STOM, SLP-2, VDAC1, BAK, STING, CASPASE-3 and CASPASE-9 were selected using the BreakingCas server (Centro Nacional de Biotecnologia, Madrid, Spain; https://bioinfogp.cnb.csic.es/tools/breakingcas/), and are shown in Table S1. Three guides were initially tested for each gene in order to select an optimal one for all experiments. Guides were ordered as pairs of TOP and BOTTOM ssDNA oligonucleotides that, after annealing, produce overhangs compatible with dissimilar Bsmb1 insertion sites present in the lentiCRISPRv2 vector series. Lentiviral plasmids containing the relevant guides were co-transfected with helper plasmids psPAX2 (Addgene Ref. 12260) and pCMV-VSV-G (Addgene, Ref. 8454) into HEK293T cells. Virus-enriched supernatants were collected 48 h post-transfection, filtered against 0.45 μM, and diluted 1:1 in fresh medium containing polybrene to a final concentration of 8 μg/ml. Infections were carried out by spinning the resulting mix onto the target cells for 1 h at 2000 rpm, 32°C. Cells were then selected for 5-7 days in the relevant antibiotic (puromycin 1 μg/ml; hygromycin 200 μg/ml; blasticidin 2 μg/ml for MEFs and 4 μg/ml for HEK-293T cells). The resulting selected cells were used for the experiments as polyclonal populations in order to avoid a possible population bias derived from the selection of random clones.

Mouse ATG16L1 depletion was carried out using guide #1 targeted to the 5’ coding region of the cDNA (nucleotides 225-244 of the RefSeq cDNA NM_001205391). Guide #2 is directed to the central area of the cDNA (nucleotides 1262-1281 of the same RefSeq file), and it is surprisingly able to deplete ATG16L1 expression almost as efficiently as guide #1. Since the target sequence (nucleotides 1262-1281) is located downstream the coding sequences for residue T300, cells harboring this guide are amenable for expression of the N-terminal fragment of ATG16L1 (1-299) to evaluate the requirement for the WD40 domain in the different assays. We followed this strategy to test the role of the WDD domain in *Bax-/-* cells.

PHB depletion caused reduced proliferation and cellular discomfort that became more obvious after 3-4 days of transduction, and long selection times resulted in the recovery of PHB expression to wild-type levels, probably due to the competitive overgrowth of non-targeted cells. Therefore, PHB depletions in all experiments were carried out acutely, aiming at a fast and incomplete depletion of the molecule that would allow functional testing with minimal side effects. To this end, cells were transduced with the CRISPR/Cas9 lentiviral vectors and, after 12-18 h, treated with the relevant selection agent for just 24 h before subjecting them to transfection or retroviral transduction (total depletion time 36-40 h). Under these conditions, PHB depletion was routinely around 80% and no obvious toxicity was detected in any of the depleted samples. The PHB2 CRISPR/Cas9 guide caused reduction in the expression levels of both PHB2 and PHB1, in line with previous evidence showing that both proteins are required to maintain their respective expression levels(*63*). Accordingly, only the CRISPR guide for PHB2 was used for depletion of both PHBs in all experiments.

siRNAs against murine VPS34, ATG14, ATG7 and ATG16L1 were mixtures of four single siRNAs targeting the same molecule of the ON-TARGETplus SMART pools collection (Dharmacon, L-063416-00, L-172696-00, L-049953-00, L-051669-01, respectively). siRNAs were transfected at a final concentration of 0.5 μM using DharmaFECT1 reagent (Dharmacon, T-2001-02), according to the manufacturer’s instructions. Control siRNAs were a mix of duplexes with no perfect match with any human or mouse gene (ON-TARGETplus, Dharmacon, D-001810-10).

### Cell lysis, Western blot and immunoprecipitation

To obtain total lysates for Western blot, cells were collected by centrifugation, washed in PBS 1x and resuspended in 2x SDS sample buffer (SB) lacking β-mercaptoethanol and bromophenol blue (100 mM Tris-HCl pH 6.8, 4% SDS and 20% de glycerol) supplemented with PMSF (1 mM) and a Protease Inhibitor Cocktail (PIC, Sigma). After vigorous vortexing, samples were boiled for 15 min and then centrifuged at 13200 rpm (16100xg) for 5 min at 4°C to remove debris. Lysates were quantified for protein concentration using the DC Protein Assay Kit (Lowry method, BioRad, 5000116). Equal amounts of protein were volume normalized with SB buffer, diluted 1:1 with 2x SDS sample buffer containing β-mercaptoethanol (10 %) and bromophenol blue (0,3 %), boiled for 10 min, spun at top speed and subjected to protein polyacrylamide gel electrophoresis plus Western blot.

For co-immunoprecipitation assays carried out in transfected HEK-293T cells, at the end of the experiment the cells were harvested, washed and lysed for 30 min (in ice, with occasional vortexing) in a buffer containing 1% Igepal CA-630 detergent (Sigma, I3021), 50 mM Tris pH 7.5, 150 mM NaCl, 5 mM EDTA, protease inhibitor cocktail (1/100 dilution; Sigma, P8430), PMSF (1 mM; Sigma, 10837091001), orthovanadate (200 nM; New England Biolabs, P0758S) and β-glycerophosphate (25 mM; Sigma, G9422). After a 10 min centrifugation step (16200xg, 4°C), the clarified lysates were diluted to a final detergent concentration of 0.2% to preserve protein-protein interactions, precleared with agarose beads (Thermo Fisher, 26150) for 30 min (4°C, rotation) and subjected to immunoprecipitation using agarose-GSH (GE-Healthcare, 17-0756-01) for 2 h (4°C, rotation). Samples were washed 2-3 times in the same immunoprecipitation buffer, resuspended in 2x Reducing Sample buffer (RSB: SB supplemented with β-mercaptoethanol (10%) and bromophenol blue (0,2%)), and boiled for 10 min. A portion of the initial lysates was saved to be used as control total lysates, whose concentrations were measured using the DC Protein Assay Kit (Lowry method, BioRad, 5000116), following the instructions provided by the manufacturer.

Co-immunoprecipitation assays in DKO MEFs reconstituted with GST-hBAK or hBAK and other components of the pathway were carried out in a derivative DKO cell line where both caspase-3 and caspase-9 were inactivated using the lentiviral v2 system (see above the CRISPR/Cas9 section). These co-immunoprecipitation experiments were challenging due to unusually high nonspecific signal provided by PHBs and STOM. To solve this difficulty, cells were first crosslinked by resuspending them in PBS containing 1% paraformaldehyde (5 min, RT) followed by a quenching step to block free aldehyde groups in Tris HCl pH 7.4, 150 mM NaCl (10 min, RT). This crosslinking step allowed extensive washing of the resulting immunoprecipitates. Cells were then lysed in the same 1% Igepal-based buffer described above and centrifuged to remove nuclei. Protein extracts were precleared (30 min, 4°C, rotation) with agarose beads (for cells expressing GST-BAK, PHBs and STOM) or magnetic beads (Dynabeads, Thermo Fisher, 11201D) coupled to a control IgG1 mouse immunoglobulin (Thermo Fisher, 14-4714-82; for cells expressing untagged BAK, PHBs and STOM). Immnoprecipitations were carried out for 4 h (4°C, rotation) using agarose-GSH (for cells expressing GST-BAK) or magnetic beads coupled to an anti-BAK antibody able to recognize the Nt region of BAK only when activated(*49*) (mouse mAb, clone G317.2, BD Biosciences, 556382, for cells expressing untagged BAK). GST-BAK immunoprecipitates were washed extensively (6 times, 5 min, 4°C, rotation) in the same 1% Igepal-based buffer. Anti-BAK precipitates were washed 2 times in RIPA buffer (5 min, 4°C, rotation) and 4 additional times in 1% Igepal (5 min, 4°C, rotation). Beads were resuspended in 2x RSB and boiled for 15 min.

Pull-down assays were carried out using GST-latch chimeric constructs purified from transfected HEK-293T. Cells were transfected with the GST-latch chimera and, after 36 h, lysed in RIPA buffer (50 mM Tris-HCl pH 7.4, 150 mM NaCl, 1% Triton X-100, 0.5% sodium deoxycholate, 0.1% SDS) containing protease inhibitor cocktail (1/100 dilution; Sigma, P8430), PMSF (1 mM; Sigma, 10837091001), orthovanadate (200 nM; New England Biolabs, P0758S) and β-glycerophosphate (25 mM; Sigma, G9422), for 30 min (ice, occasional vortexing) followed by a 10 min centrifugation step to remove debris (16200xg, 4°C). Lysates were precleared with agarose beads (30 min, 4°C, rotation) and incubated with agarose beads coupled to GSH (2 h, 4°C, rotation). Beads were then extensively washed with RIPA buffer to remove endogenous co-precipitating PHBs (6 times, 5 min at 4°C, rotation) followed by two washes in a buffer containing 0.2% Igepal CA-630, 150 mM NaCl, 50 mM Tris-HCl pH 7.5, 5 mM EDTA, for equilibration. These washed beads were used to pull-down PHBs from whole cell lysates of HEK-293T cells previously transfected with PHB1 and PHB2 expression constructs. To generate these crude lysates, cells were lysed 36 h post-transfection in the same 1% Igepal CA-630-based buffer described above. Lysates were then diluted 1:5 to reach a 0,2 % detergent concentration, and subjected to pull-down by incubation with the beads loaded with the GST-latch fusion proteins (4 h, 4°C, rotation). Beads were then washed three times in the same buffer, resuspended in 2x RSB and boiled for 10 min.

To evaluate cytochrome c release from mitochondria, cells were retrovirally transduced with constructs expressing BH3-only molecules, treated with zVAD.fmk (20 uM) 7 h later and lysed 20 h post-transduction in a buffer containing 25 mM Tris-HCl pH 7.4, 10 mM KCl, 5 mM MgCl2, 1 mM EDTA, 250 mM sucrose and 0.025% digitonin for 2 min (ice). Lysates were then centrifuged at 14000xg (4°C, 3 min) and the resulting supernatants harvested. Pellets were lysed in SB buffer. Supernatants and pellet lysates were subjected to anti-cytochrome c Western blot.

Equal amounts of protein or equal immunoprecipitation volumes were resolved by SDS-PAGE, transferred to a polyvinilidene difluoride membrane (PVDF, Millipore, IPVH00010), and probed with specific antibodies against LC3 (WB: mouse, MBL, M186-3, 1/1000), BAK (rabbit, BD Pharmingen, 556396, 1/1000; rabbit, Cell Signaling, 12105, 1/1000; rabbit, Abcam ab32371, 1/1000), BAX (rabbit, Santa Cruz Biotech, N20, sc-493), p62 (rabbit, MBL, PM045B, 1/1000), VPS34 (rabbit, Cell Signaling, 3358, 1/1000), ATG14 (rabbit, MBL, PD026B, 1/1000), ATG7 (rabbit, Cell Signaling, 2631, 1/1000), ATG16L1 (mouse, MBL, M150-3, 1/1000), PHB1 (mouse, Santa Cruz Biotech, sc-377037, 1/800), PHB2 (rabbit, Cell Signaling, 14085, 1/1000), STOM (rabbit, Abcam, ab166623, 1/1000), SLP2 (mouse, Santa Cruz Biotech, sc-376165, 1/500), VDAC1 (rabbit, Abcam, ab154856, 1/1000), BIM (rabbit, Abcam, ab15184, 1/200), Actin (mouse, Sigma AC40, A3853, 1/1000), cytochrome c (rabbit, Proteintech, 10993-1-AP, 1/1000), STING (rabbit, Proteintech, 19851-1-AP, 1/1000), phospho-STING (rabbit, Cell Signaling, 72971S, 1/500), GAPDH (mouse, Abcam, Ab8245, 1/1000), Tubulin (mouse, Sigma, T5201, 1/5000), or GFP (mouse, Covance, MMS-118P, 1/5000), Flag (mouse, Sigma, F1804, 1/1000), GST (mouse, Santa Cruz Biotech, sc-138, 1/1000), HA (rabbit, Cell Signaling, 3724, 1/1000), AU1 (rabbit, Pierce, PA1-26548, 1/1000) tags. After an additional incubation with anti-mouse (goat, Jackson Immunoresearch, 115-035-003, 1/10000) or anti-rabbit (goat, Jackson Immunoresearch, 111-035-003, 1/10000) secondary HRP-coupled antibodies, blots were developed by chemiluminescence using the ECL system (Amersham, RPN2134). When reprobed, in case of a molecular weight conflict with a previous probing, membranes were previously stripped for 10 min in a 7 M guanidinium hydrochloride solution. Quantifications were carried out with the ImageJ software.

### Immunofluorescence and microscopy

Cells were seeded onto poly-L-lysine (Sigma, P4832) coated coverslips and transfected or retrovirally transduced the next day for the indicated times. For anti-LC3 staining, coverslips were first washed with PBS and fixed with methanol (100%, −20°C, 5 min). Samples were washed, fixed in 4% paraformaldehyde for 10 min and quenched/permeabilized in PBS containing 100 mM glycine and 0.5% NP40 (RT, 30 min). For antibody staining, coverslips were blocked in PBS/3% BSA (RT, 45 min) and then stained with primary and secondary antibodies diluted in PBS/2%BSA for 1 h at RT. Primary antibodies were: anti-LC3 (1/200, 5F10, mouse mAb, NanoTools 0231-100) or anti-AU (1/1000, rabbit pAb, Thermo Fisher, PA1-26548). Secondary antibodies were AffiniPure goat anti-mouse-Cy3 (Jackson Immunoresearch, 115-165-003, 1/400) and goat anti-rabbit Alexa Fluor 488 (Molecular Probes, A11008, 1/400). Coverslips were then incubated with DAPI (1 μg/ml in PBS) for 5 min and mounted in Prolong Gold anti-fade mounting medium (Thermo Fisher, P10144). Pictures were taken in a STELLARIS 8 STED confocal microscope (Leica) running on a LAS X 4.2.0.23645 software and using the 488 nm (for GFP-LC3, GFP-ATG16L1 and mGFP, detection window 505-545 nm), 553 nm (for Cy3, mRFP and Cherry-LC3, detection window 563-630 nm), 633 nm (for Cy5, detection window 645-700 nm) and 405 nm (for DAPI, detection window 415-475 nm) laser lines. The blue signal corresponds to DAPI-stained nuclei in all pictures. Scale bars represent 10 μm in all micrographs.

For quinacrine staining, cells expressing Cherry-LC3 were seeded in coverslips, treated and, at the end of the experiment, washed twice in Krebs-Ringer buffer (25 mM Hepes pH 7.4; 125 mM NaCl; 5 mM KCl; 1 mM MgSO_4_; 0.7 mM KH_2_PO_4_; 2 mM CaCl_2_; 6 mM glucose). Staining was carried out in Krebs-Ringer buffer containing 5 μM quinacrine (Sigma, Q3251) for 15 min at 37°C. Coverslips were washed again, placed upside down on glass-bottom Wilco plates (GWSt-3522) and analyzed *in vivo* on an Okolab chamber at 37°C and 5% CO_2_ coupled to the same confocal microscope mentioned above. Pictures were taken using the 488 nm (for quinacrine, detection window 500-550 nm) and 553 nm (for Cherry-LC3, detection window 563-630 nm) laser lines. Scale bars represent 10 μm in all micrographs.

Quantification of vesicle number was carried out in all experiments by blindly counting vesicles on confocal micrographs. Colocalizations were quantified using the JACoP plugin of the Fiji program for image analysis. To quantify the extent of GFP-ATG16L1 recruitment to the mitochondrial surface in an unbiased way, pictures were adjusted for identical red and green intensity, and the percentage of green dots (GFP-ATG16L1) that generate a detectable yellow colocalization with the mRFP mitochondrial marker was then evaluated by blindly counting over the adjusted images.

### Measurement of cell death

Cell death was measured by flow cytometry using inclusive methods based on the permeability of nuclear dyes Cells were seeded in 24-well plates and treated or retrovirally transduced the next day. At the desired times, both floating and attached cells were collected for each well (pipetting and trypsin treatment, respectively), centrifuged for 10 min (6000 rpm (3300xg), 4°C) and resuspended in fresh medium containing 1 μg/ml propidium iodide (PI). DAPI (1 μg/ml) was used instead of PI to measure cell death induced by MTX, since this drug emits strong red fluorescence. Samples were incubated for 10 min and then diluted 1:5 in medium before subjecting them to flow cytometry using a FACSCanto II (Becton Dickinson) device operating with the FACSDiva 8.0.1 software. Cells were gated in the FS/SS window to include both live and dead cells and exclude small debris. Live and dead cells were then assessed for propidium iodide or DAPI fluorescence in FL3.

Cell death induced by overexpression of BAK constructs was measured in HEK-293T cells upon co-transfection with GFP to identify transfected cells. After 36 h, random pictures of the cultures were blindly taken using an inverted fluorescence microscope (Axiovert200 (Zeiss) coupled to a sCMOS Monocroma camera), and the percentage of green cells showing signs of cell death (membrane blebbing, cell shrinkage and detachment from the substrate) was scored by counting the cells showing the different phenotypes. A total of 10 fields including at least 20 cells per field were scored in all conditions.

### Electron microscopy

*Bax-/-* MEFs stably expressing GFP-LC3 were retrovirally transduced with BIM and supplemented with 30 uM zVAD.fmk 7 h post-transduction. Cells were fixed 17 h post-transduction in 4% PFA and 0.05% glutaraldehyde in 0.1 M PHEM buffer, pH 6.9, for 2 h at RT and 16 h at 4 °C. After extensive washing, cells samples were embedded in 10% gelatin. Sample blocks (< 1 mm^3^) were cryoprotected with 2.3 M sucrose at 4 °C for 16 h and rapidly frozen by immersion in liquid nitrogen. Samples were sectioned on an EM FCS cryoultramicrotome (Ultracut UCT, Leica) at −120 °C and collected in a mixture of 2.3 M sucrose and 2% methylcellulose solution (vol/vol 1:1). For immunogold labelling of GFP-LC3, thawed 90-nm-thick cryosections were incubated for 30 min at RT with a rabbit anti-GFP antibody (Chromotek, PABG1, 1:200 dilution) followed by protein A conjugated to 10 nm gold particles for 30 min. After labelling, sections were stained with a mix of 1.8% methylcellulose and 0.4% uranyl acetate and visualized at 80 kV with a JEOL JEM-1400 Flash electron microscope equipped with a One View CMOS 4 K camera (Gatan, United States).

### Measurement of secreted and intracellular ATP

Cells were seeded in 24-well plates and next day retrovirally transduced with the relevant BH3-only molecules or treated with MTX for the indicated times. Around 7-9 h after treatment, the medium was removed and replaced with 350 μl of fresh medium containing 3 μM ARL67156 (Sigma, A265). At the relevant experimental time points, a sample of the supernatants was removed for ATP measurement using the Enliten ATP assay kit (Promega, FF2000), following the instructions provided by the manufacturers. To exclude possible variations in ATP secretion due to different death kinetics occurring in the different cellular strains, we sampled the supernatants at two different times along the death process: 14-16 h post-treatment (when cell death is usually about 40-60%) and 20-26 h post-treatment at the end of the treatment (when cell death is about 85-95%). ATP secretion is displayed as the integrated, cumulative data arising from the sum of the luciferase activity units obtained at early and late times, and reflects more accurately the total amount of ATP liberated along the death process.

To measure the ATP pool that remains in the apoptotic bodies at the end of the apoptotic process, the cellular remains were harvested by mild pipetting, spun at 6000 rpm (3300xg), washed with cold PBS 1x, and the resulting pellet lysed in 20 μl of a buffer containing 1% Igepal CA-630 detergent (Sigma, I3021), 150 mM NaCl, 50 mM Tris-HCl pH 7.5 and 5 mM EDTA. The lysate was spun at 13200 rpm (16100xg) and 4°C to remove debris. The resulting supernatant was then collected to a separate tube and treated with a final concentration of 0,5% trichloroacetic acid (for 1 h in ice) for protein precipitation. Samples were spun at 13200 rpm (16100xg) and 4°C, and the resulting supernatants were neutralized by addition of 5x volumes of 20 mM Tris-HCl pH 7. Samples were measured for ATP content using the Enliten ATP assay kit (Promega, FF2000) and following the instructions provided by the manufacturer.

### Proteomics

HEK293T cells (20×10 cm plates) were transfected (jetPEI) with the mammalian expression plasmid P12 driving the expression of AU-GST-BAK-latch-WT or AU-GST-BAK-latch-5M. Cells were lysed 36 h later in a buffer containing 1% Igepal CA-630 detergent (Sigma, I3021), 150 mM NaCl, 50 mM Tris pH 7.5, 5 mM EDTA, the protease inhibitor cocktail (1/100 dilution; Sigma, P8430), PMSF (1 mM; Sigma, 10837091001), orthovanadate (200 nM; New England Biolabs, P0758S) and β-glycerophosphate (25 mM; Sigma, G9422). Lysates were centrifuged at 13200 rpm (16100xg; 5 min, 4°C) and the resulting supernatants diluted 1:5 in detergent-free lysis buffer to reach a total detergent concentration of 0.2%. Samples were precleared with agarose beads for 1 h (4°C, rotation) and then incubated with agarose-GSH beads for 3 h (4°C, rotation). Precipitates were washed 2x with immunoprecipitation buffer (0.2% detergent), resuspended in 2x RSB and boiled. Samples were resolved by PAGE and the gel was silver stained. Broad regions showing more intense bands in the AU-GST-BAK-latch-WT sample compared to the AU-GST-BAK-latch-5M control were excised and processed for proteomics identification.

Sample processing for mass spectrometry was carried out as previously described(*54*). Briefly, silver-stained gel plugs were destained with a solution containing 7.5 mM potassium ferricyanide and 25 mM sodium thiosulfate, rinsed in water and dehydrated with 100% acetonitrile. Plugs were then DTT reduced and alkylated with iodoacetamide. Modified porcine trypsin (Promega, Madison, WI; 6 ng/μl in 20 mM ammonium bicarbonate) was added before incubation at 37°C for 18 h. Tryptic peptides were dried in a speed vacuum system, desalted by using C18-home-made columns and analyzed by reversed-phased LC-MS/MS using a nanoAcquity UPLC (Waters Corp., Milford, MA) coupled to a LTQ-Orbitrap Velos (Thermo-Fisher, San Jose, CA). Separations were done in a BEH 1.7 μm, 130Å, 75 μm x 100 mm C18 column (Waters Corp., Milford, MA) at a 400 nL/min flow rate. Injected samples were trapped on a Symmetry, 5 μm particle size, 180 μm x 20 mm C18 column (Waters Corp., Milford, MA). Peptides were eluted using a 30 min gradient from 3 to 35% B (0.5% formic acid in acetonitrile). The LTQ-Orbitrap Velos was operated in a data-dependent MS/MS mode using Xcalibur (Thermo-Fisher, San Jose, CA). Survey scans were acquired in the mass range 400-1,600 m/z, with 30,000 resolution at m/z 400 and lock mass option enabled for the 445.120025 ion. The 20 most intense peaks having ≥ 2 charge state and above 500 intensity threshold were selected in the ion trap for fragmentation by collision-induced dissociation. MASCOT [v 2.3] and Sequest HT [v 1.3] search algorithms were used for searching the acquired MS/MS spectra using Thermo Scientific Proteome Discoverer software (v. 1.4.1.14) against a database of human sequences (Uniprot) with common contaminants. Search parameters were as follows: fully-tryptic digestion with up to two missed cleavages, 10 ppm and 0.8 Da mass tolerances for precursor and product ions, respectively, oxidation of methionine was established as variable modification and carbamidomethylation of cysteine as fixed modification. 1% false discovery rate using Percolator was used for peptide validation.

Peptides identified and the corresponding proteins (a total of 661) were downloaded on an excel spreadsheet and, ordered according to a decreasing difference in the number of peptides identified in the latch-WT sample compared to the latch-5M (Source Data 1). From this collection, 239 proteins were selected as having at least one more peptide present in the latch-WT sample compared to the control (Source Data 2), 35 of which satisfied the arbitrary threshold of being overrepresented in the WT sample by more than 4 peptides (highlighted candidates in Source Data 2). These were selected for further analysis. The mass spectrometry proteomics data have been deposited to the ProteomeXchange Consortium via the PRIDE partner repository with the dataset identifier PXD045985 and 10.6019/PXD045985.

To identify proteins commonly present in the BAK latch proteomics list that we identified in this work and the ATG16L1-WDD proteomics approach that we previously published(*54*), we resorted to a Venn diagram (Venny 2.1 sever; https://bioinfogp.cnb.csic.es/tools/venny/). We confronted a crude, unfiltered list of ATG16L1-WDD hits including 1646 candidates (curated as Uniprot reviewed references) with the BAK latch results shown in Source Data 2. The Venn diagram indicated that a total of 117 proteins were commonly present in both lists, including PHB1 and STOM (Source Data 3).

### IL-1β release by bone marrow-derived macrophages (BMDMs)

Wild-type C57BL6/J mice were euthanized using a CO_2_ chamber and dissected to isolate bone marrow cells by flushing the tibias and femurs with a 23G syringe. Red cells were lysed with an ACK Lysing Buffer (Life Technologies, A1049201), and the resulting suspension was plated at 1-2×10^6^ cells/ml in complete medium containing 20% FCS and 20 ng/ml M-CSF (BioLegend, 574804). Cells were incubated for 6 days with a medium replacement at day 3. At the end of this differentiation procedure, purity was 100% as measured by two-color flow cytometry for double surface expression of CD11b (rat anti-mouse CD11b-BB515, Becton Dickinson, 564454) and F4/80 (rat anti-mouse F4/80-Alexa Fluor 647, Becton Dickinson, 565853); data not shown. Cells were detached with Accutase (Sigma, A6964) and plated in 24-well plates at 3-3.5×10^5^ cells per well. Next day, cells were treated with LPS (ultrapure, Invivogen, Tlrl-3pelps; 100 ng/ml) for 4 h, and then with a 1:5 proportion of the relevant apoptotic bodies for 6 h or with ATP (5 mM) for 30 min as a positive control. Supernatants were harvested for IL-1β measurement using a specific ELISA (Thermo Fisher, 88-7621-22). Apoptotic cells were generated by retroviral transduction or MTX treatment during 20-24 h when cell death approaches its maximum. ARL67156 (Sigma, A265; 3 μM) was added 7 h post-treatment to prevent ATP degradation. At the end of the procedure, apoptotic bodies were harvested by soft pipetting, spun at 6000 rpm (3300xg; 4°C, 10 min), resuspended in 350 ml of their own supernatant (which contains secreted ATP), and added to the macrophage cultures pre-stimulated with LPS. Control samples were the supernatants of live cells transduced with irrelevant vector or treated with vehicle. To test the effect of P2×7 channel inhibitors, macrophages were pre-incubated with the indicated doses for 45 min and the apoptotic bodies were added to the BMDM cultures in medium containing the same dose of the inhibitor. Concentrations were calculated using a standard curve established according to the manufacturer’s instructions.

### Animal and ethical issues

Wild-type C57BL6/J mice for BMDM isolation were maintained in a standard animal facility using ventilated racks under controlled temperature (23°C), humidity (50%) and light/dark cycle (12 h/12 h) conditions, strictly following European Union regulations. Maintenance conditions and procedures were evaluated and approved by the Ethics and Animal Wellbeing committees of both the research center (Centro de Biologia Molecular Severo Ochoa, CBMSO) and the hosting institution (Consejo Superior de Investigacones Cientificas, CSIC).

### Statistics

The displayed graphs show means -/+ standard deviations obtained from the indicated number of experimental points (n), and were created using GraphPad Prism 8. Statistical significance (*P*-values) was calculated in all cases using two-tailed Student’s *t*-tests.

## ACKNOWLEDGEMENTS

We thank Drs. Mario Mairhofer (University of Applied Sciences, Austria), Joaquín Mádrenas (UCLA Medical School, Los Angeles, USA), Dirk Goerlich (Max Plack Institute, Gottingen, Germany), Craig Thompson (Sloan Kettering Institute, New York, USA) and Bert Vogelstein (Johns Hopkins School of Medicine, Baltimore, Maryland, USA) for the STOM, SLP2, EXP4, BCLXL and PUMA cDNAs, respectively. We also thank Dr. Javier Traba for advice on macrophage protocols, Dr. Emilio Lecona and Rodrigo Martín for help and reagents used in the co-precipitation studies, and CBMSO facilities and personnel for support. This work was funded by grants SAF2017-88390-R and PID2020-114699RB-100 from the Ministerio de Ciencia y Tecnología (Spanish Government), IDEAS18093PIME from the Asociación Española Contra el Cáncer (AECC) and SA042P17 from the Junta de Castilla y León local government. Additional funding comes from the of Excellence Severo Ochoa Grant CEX2021-001154-S from MCIN/AEI /10.13039/501100011033. The CBM receives institutional support by Fundación Ramon Areces. E.T. and J.B.L. are recipients of predoctoral fellowships PRE2018-085243 and PRE2021-097461, respectively, from the Formación de Personal Investigador (FPI) program of the Spanish Ministry of Science and Technology (MCIU/AEI/10.13039/501100011033; Spanish Government and FSE+). F.X.P. holds a tenured position at CSIC.

## AUTHOR CONTRIBUTIONS

E.T.S, R.V., J.B.L., M.L., C.R.B., C.M.J. and D.O.S. performed the experiments. F.X.P. conceived and designed the study, and wrote the manuscript.

## DECLARTION OF INTERESTS

Authors declare that they have no competing interests.

## DATA AVAILABILITY STATEMENT

Data and materials are freely available upon request to the corresponding author. The mass spectrometry proteomics data have been deposited to the ProteomeXchange Consortium via the PRIDE partner repository with the dataset identifier PXD045985 and 10.6019/PXD045985.

