## Supplementary Figures and Tables for "An unconventional autophagic pathway that inhibits ATP secretion during apoptotic cell death"

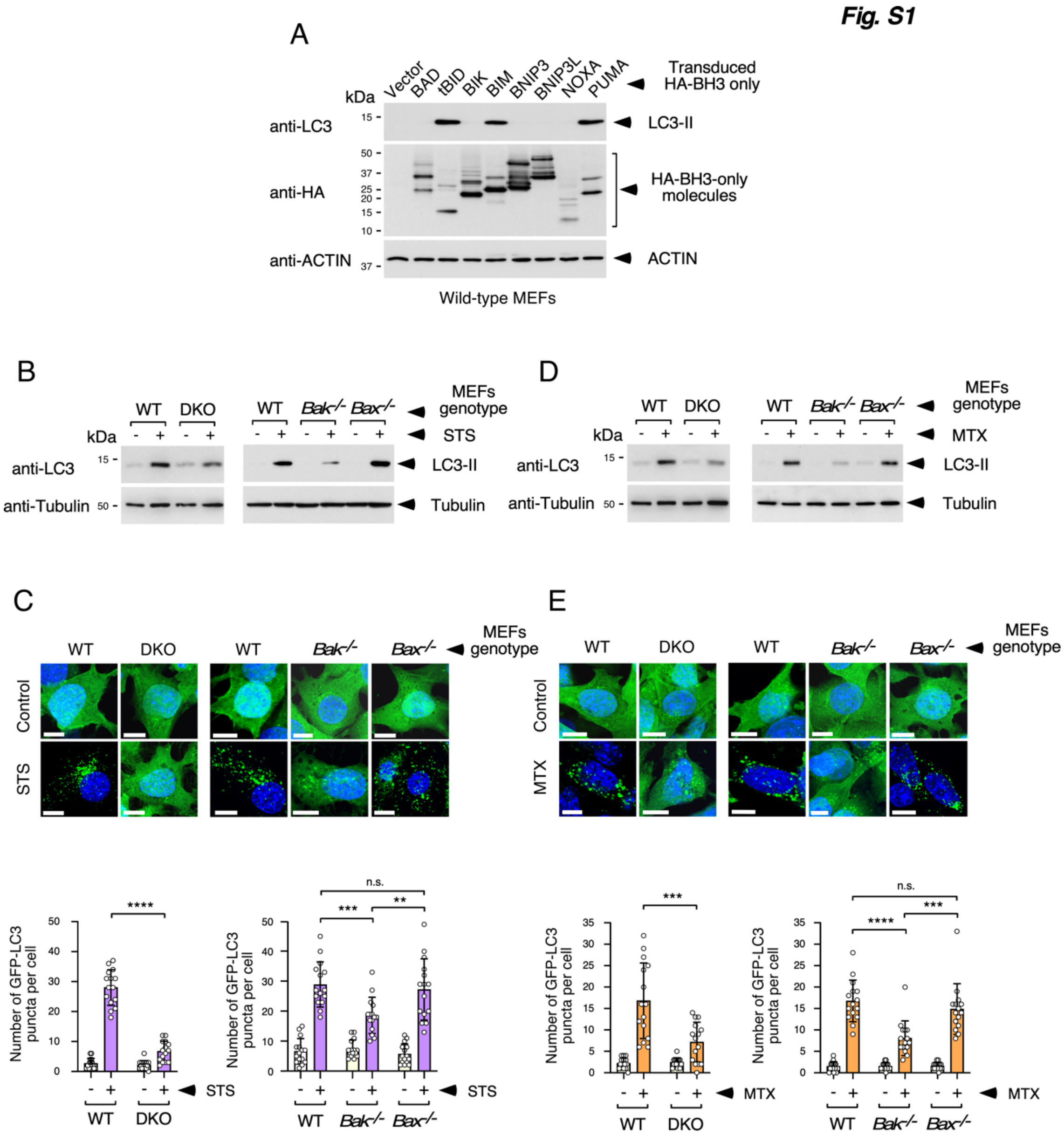

**Suppl. Fig. 1. LC3 lipidation induced by different BH3-only molecules, and role of BAK in the autophagic response triggered by cytotoxic drugs. (A)** LC3 lipidation induced by expression of BH3-only molecules. Wild-type MEFs were transduced with retroviral constructs expressing the indicated HA-tagged BH3-only molecules and supplemented with 25 μM zVAD.fmk 7 h later. Cells were lysed 22 h post-transduction for Western blot with the indicated antibodies. (**B, D)** Staurosporine (STS) and mitoxantrone (MTX) induce BAK-dependent LC3 lipidation. The indicated MEFs were treated with STS (2 μM, 16 h, **B**) or MTX (2 μM, 16 h, **D**) in the presence of 25 μM zVAD.fmk, and lysed for Western blot. **(C, E)** STS and MTX induce BAK-dependent GFP-LC3 activation. MEFs expressing GFP-LC3 were treated with STS (2 μM, 16 h, **C**) or MTX (2 μM, 16 h, **E**) and fixed for microscopy. Shown are representative confocal pictures (top panels) and quantification of the number of GFP-LC3 puncta per cell (bottom panels). Graphs represent mean values -/+ s.d. (n = 15 cells; n.s. $P>0.05$, **$P<0.01$, ***$P<0.001$, ****$P<0.0001$ Student's *t*-test).

*Fig. S2*

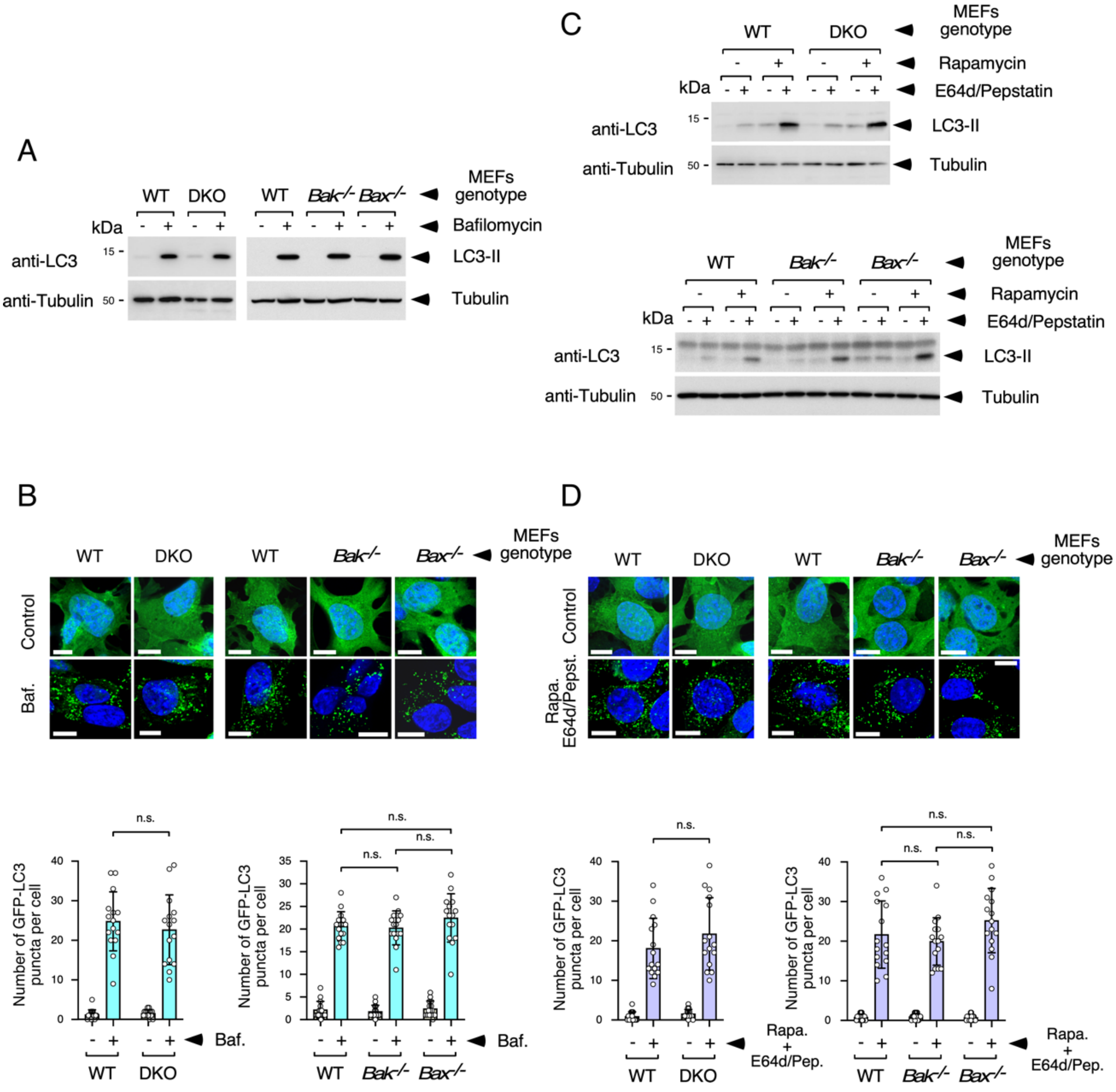
A
WT
DKO
WT
Bak-/-
Bax-/-
MEFs genotype
kDa
Bafilomycin
anti-LC3
15
LC3-II
anti-Tubulin
50
Tubulin
B
Control
Baf.
Number of GFP-LC3 puncta per cell
n.s.
Baf.
C
Rapamycin
E64d/Pepstatin
D
Rapa. E64d/Pepst.
Rapa. + E64d/Pep.

**Suppl. Fig. 2. Unaltered basal and canonical autophagy in DKO and *Bak-/-* MEFs. (A, C)** LC3 lipidation induced by basal and canonical autophagy is unaffected in MEFs lacking BAK and/or BAX. The indicated MEFs were treated with bafilomycin (for basal autophagy, 80 nM, **A**) or rapamycin (2 μM) plus E64d/pepstatin (for canonical autophagy, 10 μg/ml each, **C**) and lysed 8 h later for Western blot. **(B, D)** GFP-LC3 activation induced by basal and canonical autophagy is unaffected in MEFs lacking BAK and/or BAX. MEFs expressing GFP-LC3 were treated with (80 nM, **B**) or rapamycin (2 μM) plus E64d/pepstatin (10 μg/ml each, **D**) and fixed 8 h later for microscopy. Shown are representative confocal pictures (top panels) and quantification of the number of GFP-LC3 puncta per cell (bottom panels). Graphs represent mean values -/+ s.d. (n = 15 cells; n.s. $P>0.05$ Student's *t*-test).

***Fig. S3***

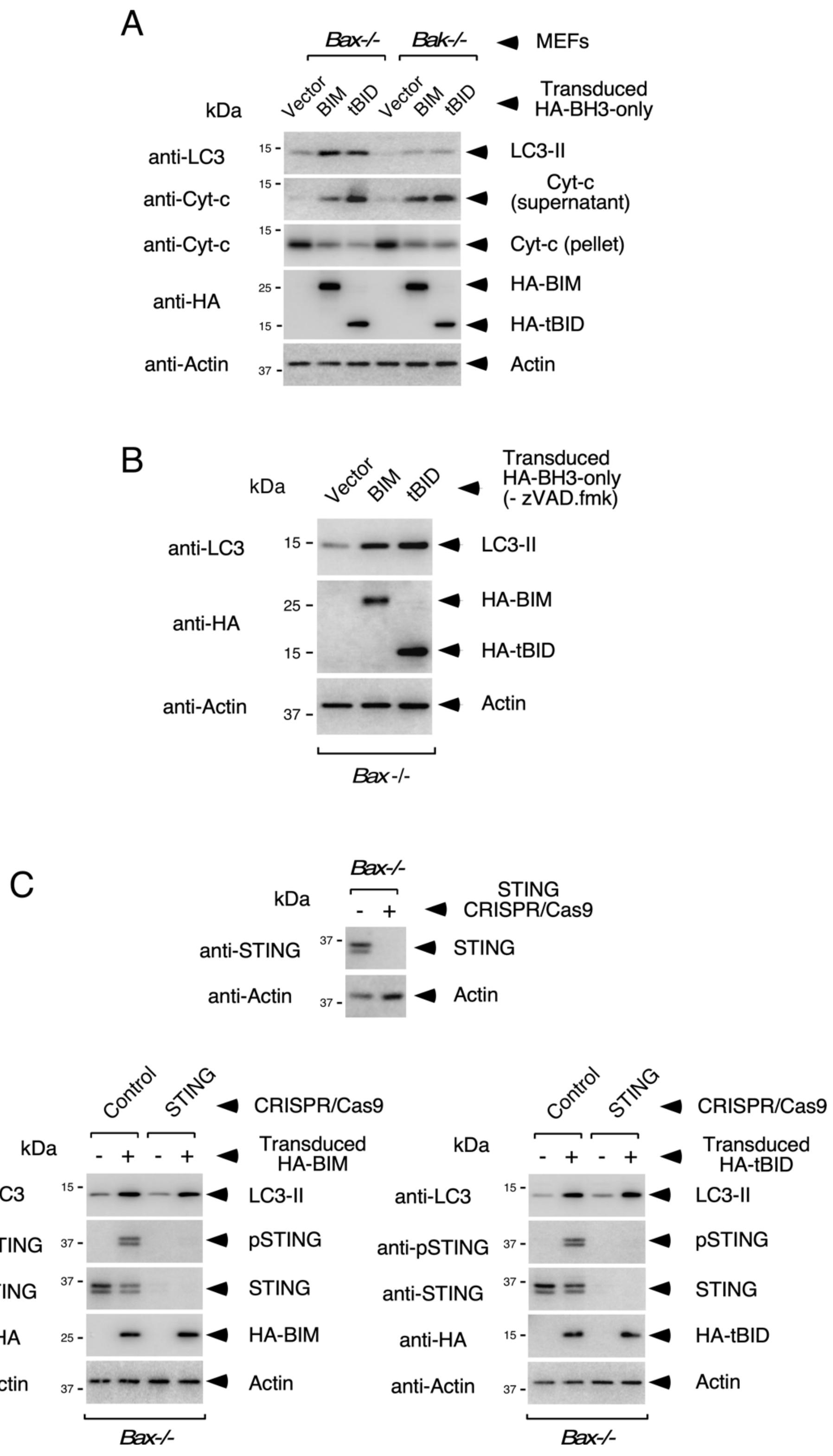
A
Bax-/-
Bak-/-
MEFs
kDa
Vector
BIM
tBID
Vector
BIM
tBID
Transduced HA-BH3-only
anti-LC3
LC3-II
anti-Cyt-c
Cyt-c (supernatant)
anti-Cyt-c
Cyt-c (pellet)
anti-HA
HA-BIM
HA-tBID
anti-Actin
Actin
15
15
15
25
15
37
B
kDa
Vector
BIM
tBID
Transduced HA-BH3-only (- zVAD.fmk)
anti-LC3
LC3-II
anti-HA
HA-BIM
HA-tBID
anti-Actin
Actin
15
25
15
37
Bax -/-
C
Bax-/-
kDa
STING CRISPR/Cas9
anti-STING
STING
anti-Actin
Actin
37
37
Control
STING
CRISPR/Cas9
kDa
Transduced HA-BIM
anti-LC3
LC3-II
anti-pSTING
pSTING
anti-STING
STING
anti-HA
HA-BIM
anti-Actin
Actin
Bax-/-
Control
STING
CRISPR/Cas9
kDa
Transduced HA-tBID
anti-LC3
LC3-II
anti-pSTING
pSTING
anti-STING
STING
anti-HA
HA-tBID
anti-Actin
Actin
Bax-/-

**Suppl. Fig. 3. The autophagic response triggered by BH3-only molecules is unrelated to mitochondrial permeabilization and does not require the STING pathway. (A)** Comparable cytochrome c liberation induced by BIM and tBID in *Bax-/-* and *Bak-/-* MEFs. The indicated cells were transduced with retroviruses expressing the shown BH3-only molecules and treated with 25 μM zVAD.fmk 7 h later. Cells were lysed 20 h post-transduction for Western blot against the shown molecules. Samples for cytochrome c release were processed separately according to a specific protocol. **(B)** BIM and tBID induce LC3 lipidation in the absence of zVAD.fmk. *Bax-/-* MEFs were transduced with retroviruses expressing the shown BH3-only molecules and lysed 17 h post-transduction for Western blot against the shown molecules. Cells were not completely dismantled at this early time point. **(C)** Absence of STING does not blunt the autophagic response induced by BIM and tBID. *Bax-/-* MEFs were transduced with a CRISPR/Cas9 construct targeting STING and lysed for Western blot to assess STING expression (top panel). Cells were then transduced with the shown BH3-only molecules, treated with 25 μM zVAD.fmk 7 h later and lysed 20 h post-transduction for Western blot against the shown molecules (bottom panels).

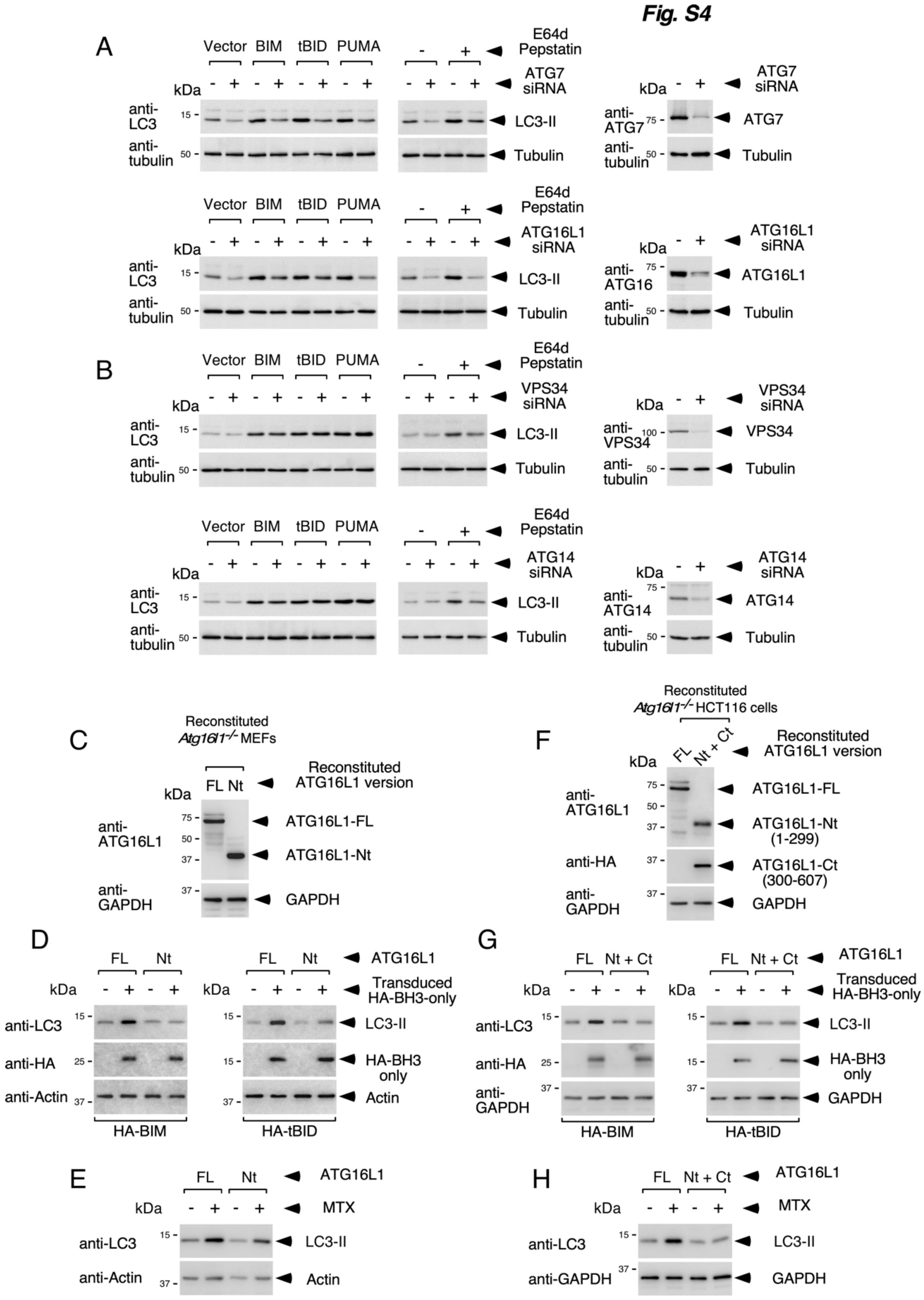
Fig. S4
A
B
C
D
E
F
G
H
Vector
BIM
tBID
PUMA
E64d
Pepstatin
ATG7
siRNA
ATG16L1
siRNA
VPS34
siRNA
ATG14
siRNA
kDa
anti-LC3
anti-tubulin
anti-ATG7
anti-ATG16
anti-VPS34
anti-ATG14
LC3-II
Tubulin
ATG7
ATG16L1
VPS34
ATG14
Reconstituted Atg16l1-/- MEFs
Reconstituted Atg16l1-/- HCT116 cells
Reconstituted ATG16L1 version
FL
Nt
Nt + Ct
anti-ATG16L1
anti-HA
anti-GAPDH
anti-Actin
ATG16L1-FL
ATG16L1-Nt
ATG16L1-Nt (1-299)
ATG16L1-Ct (300-607)
GAPDH
Actin
Transduced HA-BH3-only
HA-BH3 only
HA-BIM
HA-tBID
MTX

**Suppl. Fig. 4. Unconventional nature of the autophagic response induced during cell death. (A, B)** *Bax-/-* MEFs were transfected with siRNAs targeting the shown ATGs and, 48 h later, either transduced with the indicated BH3-only molecules and treated with zVAD.fmk 7 h later, or treated with E64d/pepstatin (10 μg/ml each). Cells were lysed 20 h and 8 h later, respectively, for Western blot against the indicated molecules. Separate samples were lysed 48 h after siRNA transfection to show ATG depletion (right panels). **(C, D, E)** *Atg16l1-/-* MEFs expressing ATG16L1-ΔWD40 (Nt) show defective LC3 activation during apoptosis. **(C)** *Atg16l1-/-* MEFs reconstituted with FL or Nt (1-299) versions of ATG16L1 were lysed for Western blot against the shown molecules. **(D, E)** MEFs were transduced with the indicated BH3-only proteins and supplemented with 25 μM zVAD.fmk 7 h later **(D)**, or treated with 2 μM MTX in the presence of 25 μM zVAD.fmk **(E)**, and lysed 20 h and 14 h later, respectively, for Western blot. **(F, G, H)** *Atg16l1-/-* HCT116 cells expressing separate Nt and Ct domains of ATG16L1 show defective LC3 activation during apoptosis. **(F)** *Atg16l1-/-* HCT116 cells reconstituted with FL or Nt (1-299) + Ct (300-607) domains of ATG16L1 were lysed for Western blot. **(G, H)** Cells were transduced with the shown BH3-only activators and supplemented with 25 μM zVAD.fmk 7 h later **(G)**, or treated with 2 μM MTX in the presence of 25 μM zVAD.fmk **(H)**, and lysed 20 h or 14 h later, respectively, for Western blot.

**Fig. S5**

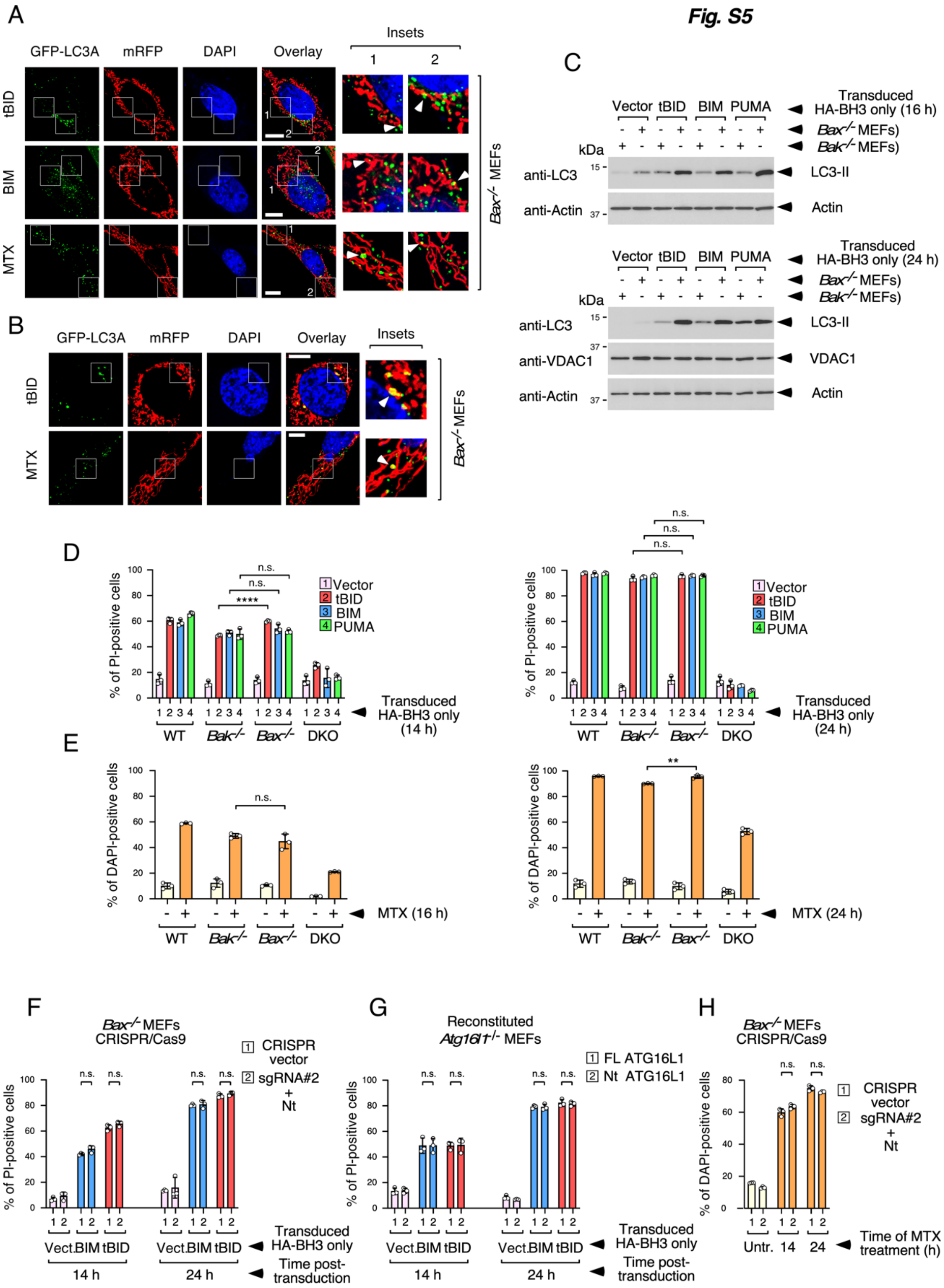

A
GFP-LC3A
mRFP
DAPI
Overlay
Insets
1
2
tBID
BIM
MTX
Bax-/- MEFs
B
GFP-LC3A
mRFP
DAPI
Overlay
Insets
tBID
MTX
Bax-/- MEFs
C
Vector tBID BIM PUMA
Transduced HA-BH3 only (16 h)
Bax-/- MEFs)
Bak-/- MEFs)
kDa
anti-LC3
anti-Actin
LC3-II
Actin
Vector tBID BIM PUMA
Transduced HA-BH3 only (24 h)
Bax-/- MEFs)
Bak-/- MEFs)
kDa
anti-LC3
anti-VDAC1
anti-Actin
LC3-II
VDAC1
Actin
D
% of PI-positive cells
Vector
tBID
BIM
PUMA
n.s.
****
WT
Bak-/-
Bax-/-
DKO
Transduced HA-BH3 only (14 h)
Transduced HA-BH3 only (24 h)
E
% of DAPI-positive cells
n.s.
**
MTX (16 h)
MTX (24 h)
WT
Bak-/-
Bax-/-
DKO
F
Bax-/- MEFs CRISPR/Cas9
1 CRISPR vector
2 sgRNA#2 + Nt
% of PI-positive cells
Vect. BIM tBID
14 h
24 h
Transduced HA-BH3 only
Time post-transduction
G
Reconstituted Atg16l1-/- MEFs
1 FL ATG16L1
2 Nt ATG16L1
% of PI-positive cells
Vect. BIM tBID
14 h
24 h
Transduced HA-BH3 only
Time post-transduction
H
Bax-/- MEFs CRISPR/Cas9
1 CRISPR vector
2 sgRNA#2 + Nt
% of DAPI-positive cells
Untr. 14 24
Time of MTX treatment (h)

**Suppl. Fig. 5. The autophagic response does not involve mitophagy nor it modulates the course of cell death. (A, B, C)** Uncommon mitophagy in *Bax*-/- MEFs in response to BH3-only proteins or MTX. **(A)** *Bax-/-* MEFs expressing GFP-LC3 and mitochondrial RFP (mRFP) were transduced with the shown BH3-only molecules for 17 h (with addition of 25 μM zVAD.fmk 7 h post-transduction), or treated with 2 μM MTX for 14 h in the presence of 25 μM zVAD.fmk, and processed for microscopy. Shown are representative confocal pictures. White arrows indicate close apposition events between GFP-LC3-positive structures and mRFP to the point of producing a yellow signal. **(B)** Confocal pictures taken from the same samples as in **A** displaying rare mitophagy events where GFP-LC3 tightly surrounds and engulfs the mRFP mitochondrial signal (white arrows). **(C)** *Bax-/-* or *Bak-/-* MEFs were transduced with the shown BH3-only proteins, supplemented 7 h later with 25 μM zVAD.fmk, and lysed 16 h (top) or 24 h (bottom) after transduction for Western blot. **(D, E)** *Bak*-/- and *Bax*-/- MEFs die to a similar extent in response to BH3-only proteins or MTX as tested by flow cytometry. **(D)** MEFs were transduced with the shown BH3-only molecules and processed 14 h (left) or 24 h (right) later to measure cell death. **(E)** MEFs were treated with MTX (2 μM) and processed 16 h (left) or 24 h (right) later. **(F, G, H)** Absence of the ATG16L1 WD40 domain does not influence the course of apoptosis measured by flow cytometry. **(F)** The indicated *Bax-/-* MEFs engineered for ATG16L1 expression were transduced with BH3-only molecules and processed 14 h or 24 h later to measure cell death. **(G)** *Atg16l1-/-* MEFs reconstituted with FL or Nt ATG16L1 were transduced with BH3-only activators and processed 14 h or 24 h later. **(H)** The indicated *Bax-/-* MEFs engineered for ATG16L1 expression were treated with MTX (2 μM) and processed 14 h or 24 h later. Graphs show mean values -/+ s.d. of the percentage of propidium-iodide positive cells obtained from triplicate experimental points (n = 3; n.s. $P$>0.05, **$P$<0.01, ****$P$<0.0001 Student's *t*-test).

*Fig. S6*

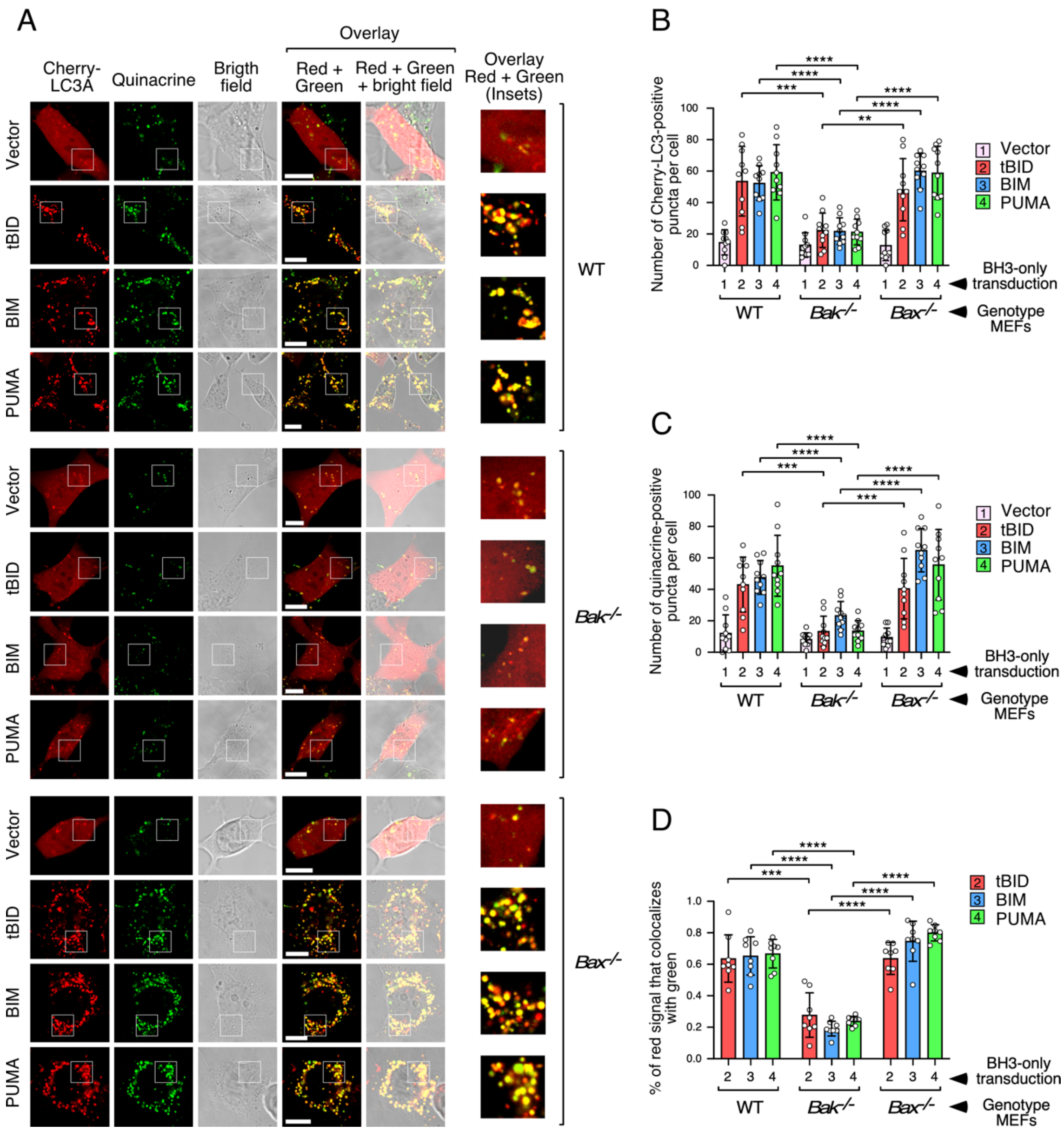

A
Cherry-LC3A
Quinacrine
Brigth field
Overlay
Red + Green
Red + Green + bright field
Overlay Red + Green (Insets)
Vector
tBID
BIM
PUMA
WT
Bak-/-
Bax-/-
B
Number of Cherry-LC3-positive puncta per cell
C
Number of quinacrine-positive puncta per cell
D
% of red signal that colocalizes with green
1 Vector
2 tBID
3 BIM
4 PUMA
BH3-only transduction
Genotype MEFs

**Suppl. Fig. 6. The LC3-positive vesicles generated during apoptosis induced by BH3-only molecules in WT and *Bax-/-* MEFs are stained with the ATP-sensitive probe quinacrine. (A)** The indicated cells stably expressing Cherry-LC3 were retrovirally transduced with BH3-only molecules, treated with 25 μM zVAD.fmk 7 h later and processed for quinacrine staining 17 h post-transduction. Samples were analyzed *in vivo* by confocal microscopy. Shown are representative confocal pictures. **(B, C, D)** Quantification of the phenotypes shown in **A**. Graphs display mean values -/+ s.d. of the number of Cherry-LC3-positive puncta per cell **(B)**, the number of quinacrine-positive puncta per cell **(C)** (in both cases, n = 10 cells; ***P*<0.01, ****P*<0.001, *****P*<0.0001 Student's *t*-test), and the percentage of Cherry-LC3-positive signal (red) that colocalizes with the quinacrine-positive signal (green) **(D)** (n = 8 cells; ****P*<0.001, *****P*<0.0001 Student's *t*-test), in the different conditions.

***Fig. S7***

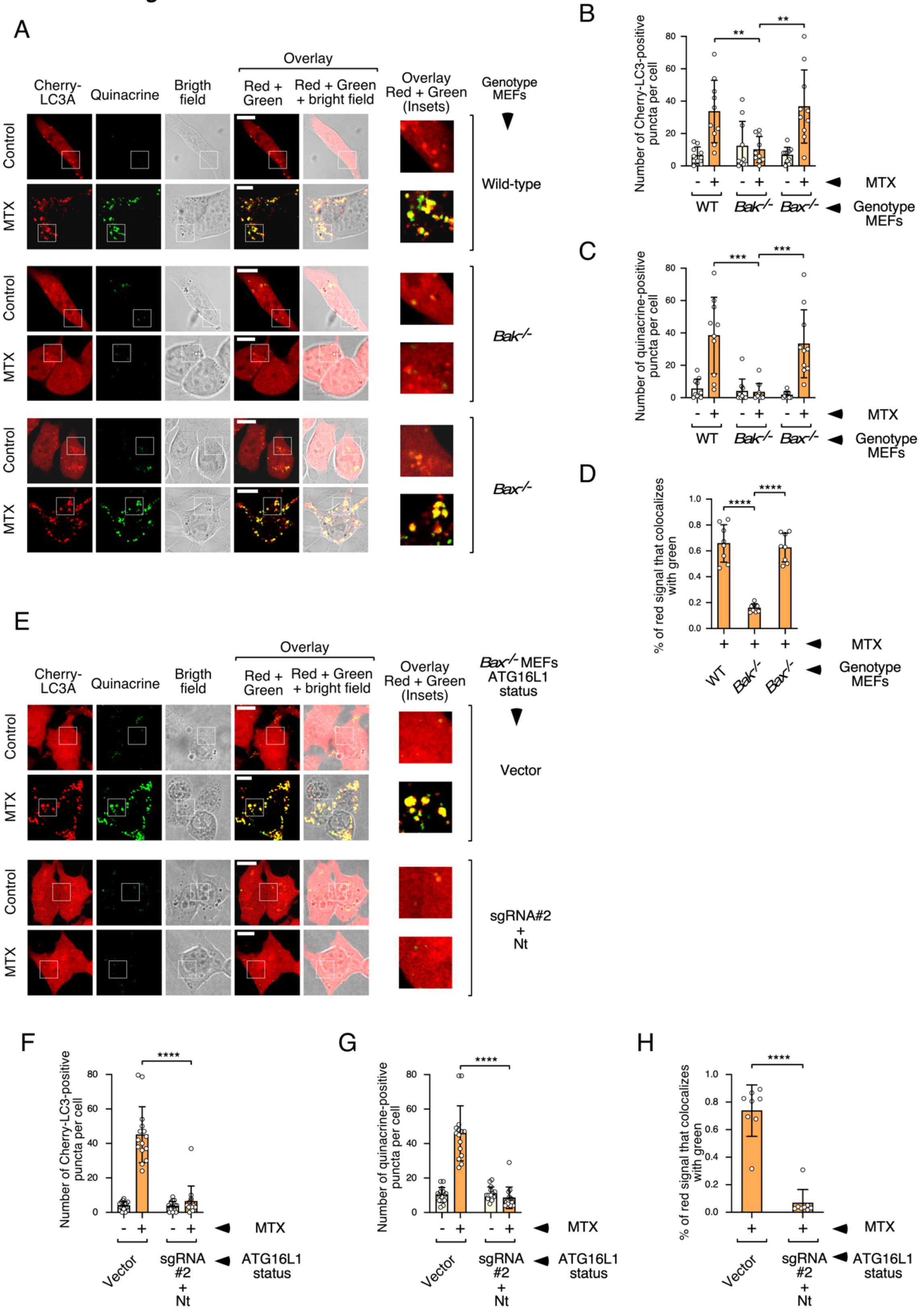
A
Cherry-LC3A
Quinacrine
Brigth field
Overlay
Red + Green
Red + Green + bright field
Overlay Red + Green (Insets)
Genotype MEFs
Control
MTX
Wild-type
Bak-/-
Bax-/-
B
Number of Cherry-LC3-positive puncta per cell
MTX
Genotype MEFs
WT
C
Number of quinacrine-positive puncta per cell
D
% of red signal that colocalizes with green
E
Bax-/- MEFs ATG16L1 status
Vector
sgRNA#2 + Nt
F
G
H
ATG16L1 status

**Suppl. Fig. 7. The LC3-positive vesicles generated during apoptosis induced by MTX treatment are stained with the ATP-sensitive probe quinacrine. (A)** The LC3-positive vesicles generated by MTX in BAK-expressing cells are stained with quinacrine. Cells stably expressing Cherry-LC3 were treated with MTX (2 μM) in the presence of 25 μM zVAD.fmk and processed for quinacrine staining 12 h later. Samples were analyzed *in vivo* by confocal microscopy. Shown are representative confocal pictures. **(B, C, D)** Quantification of the phenotypes shown in **A**. Graphs display mean values -/+ s.d. of the number of Cherry-LC3-positive puncta per cell **(B)**, the number of quinacrine-positive puncta per cell **(C)** (in both cases, n = 10 cells; **$P<0.01$, ***$P<0.001$, ****$P<0.0001$ Student's *t*-test), and the percentage of Cherry-LC3-positive signal (red) that colocalizes with the quinacrine-positive signal (green) **(D)** (n = 8 cells; ***$P<0.001$, ****$P<0.0001$ Student's *t*-test), in the different conditions. **(E)** Engineered *Bax-/-* cells expressing the Nt domain of ATG16L1 display reduced number of quinacrine-positive vesicles in response to MTX. Cells stably expressing Cherry-LC3 were treated with MTX (2 μM) in the presence of 25 μM zVAD.fmk and processed for quinacrine staining 12 h later. Samples were analyzed *in vivo* by confocal microscopy. Shown are representative confocal pictures. **(F, G, H)** Quantification of the phenotypes shown in **E**. Graphs display mean values -/+ s.d. of the number of Cherry-LC3-positive puncta per cell **(F)**, the number of quinacrine-positive puncta per cell **(G)** (in both cases, n = 15 cells; **$P<0.01$, ***$P<0.001$, ****$P<0.0001$ Student's *t*-test), and the percentage of Cherry-LC3-positive signal (red) that colocalizes with the quinacrine-positive signal (green) **(H)** (n = 8 cells; ***$P<0.001$, ****$P<0.0001$ Student's *t*-test), in the different conditions.

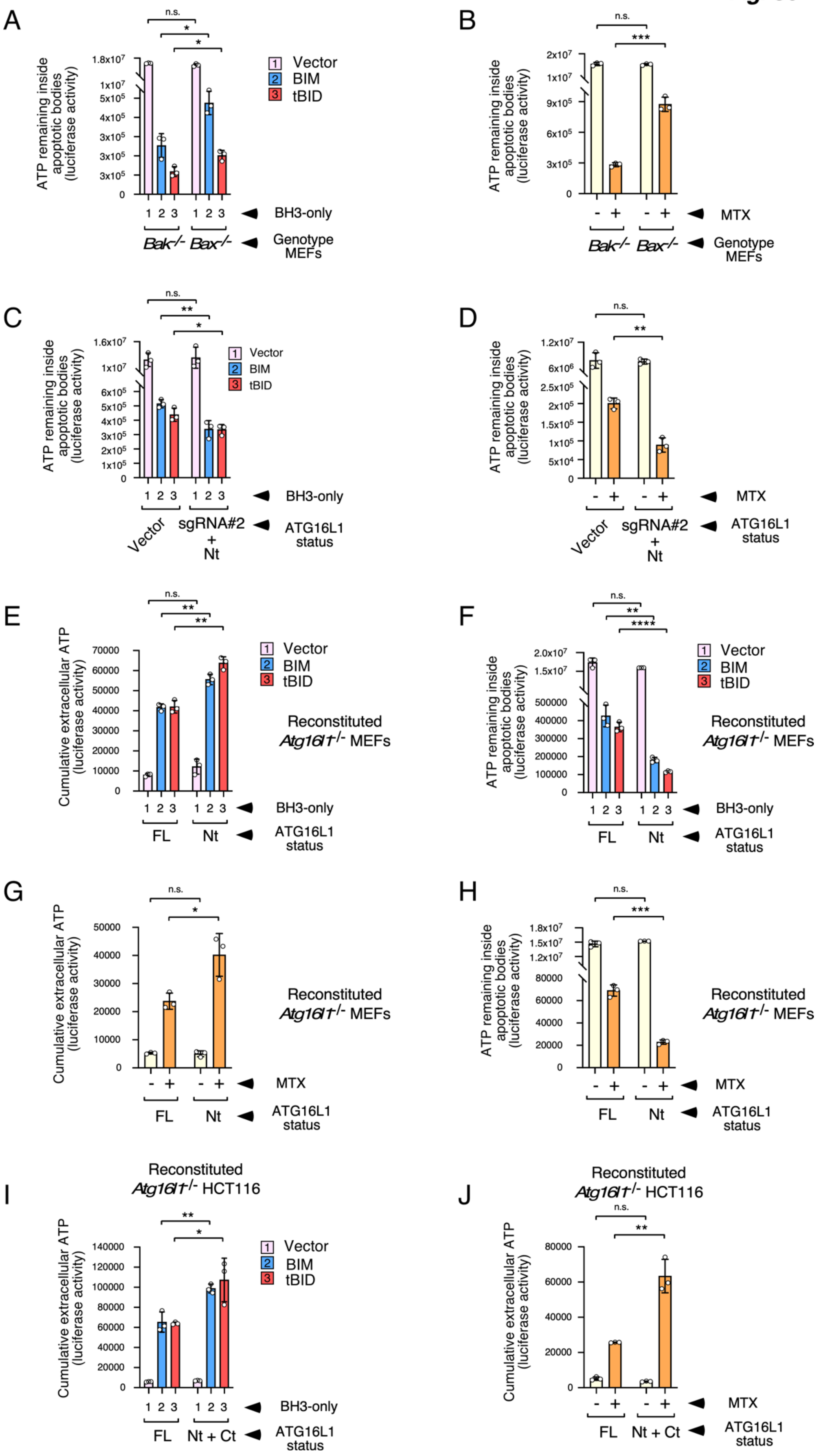
A
B
C
D
E
F
G
H
I
J
ATP remaining inside apoptotic bodies (luciferase activity)
Cumulative extracellular ATP (luciferase activity)
Vector
BIM
tBID
BH3-only
MTX
Genotype MEFs
ATG16L1 status
Bak-/-
Bax-/-
Vector
sgRNA#2 + Nt
FL
Nt
Nt + Ct
Reconstituted Atg16l1-/- MEFs
Reconstituted Atg16l1-/- HCT116
n.s.

**Suppl. Fig. 8. The unconventional autophagic response triggered during apoptosis inhibits ATP release.** All graphs show mean values of luciferase activity units -/+ s.d. from triplicate experimental points (n = 3; n.s. $P>0.05$, $^{**}P<0.01$, $^{***}P<0.001$, $^{****}P<0.0001$ Student's *t*-test). **(A, B)** Reduced ATP levels remaining in the apoptotic bodies of *Bak-/-* cells transduced with BH3-only molecules (22 h; **A**) or treated with MTX (2 μM; 24 h; **B**) compared to their *Bax-/-* counterparts. **(C, D)** Reduced ATP levels remaining in the apoptotic bodies of engineered *Bak-/-* MEFs lacking the WDD domain in response to BH3-only molecules (20 h; **C**) or MTX (2 μM; 22 h; **D**). **(E, G)** Increased ATP release caused by BH3-only molecules (transduced for 22 h; **E**) or MTX (treated for 24 h; 2 μM; **G**) in cells lacking the WDD, using as a model system *Atg16l1-/-* MEFs reconstituted with FL or Nt ATG16L1. ATP was measured at 14 and 22h (BH3-only) and 16 and 24 h (MTX), and the graphs display cumulative data resulting from the sum of the luciferase activity units obtained at both time points. **(F, H)** Reduced ATP levels remaining in apoptotic bodies of cells lacking the WDD in response to BH3-only molecules (22 h; **F**) or MTX (2 μM; 24 h; **H**) using the same cells as in **E** and **G**. **(I, J)** Increased ATP release caused by BH3-only molecules (22 h; **I**) or MTX (26 h; 2 μM; **J**) in cells lacking WDD functionality using as a model system *Atg16l1-/-* HCT116 cells expressing ATG16L1 FL or separated Nt + Ct domains. ATP was measured at 14 and 22h (BH3-only) and 18 and 26 h (MTX), and the graphs display cumulative data as in **E** and **G**.

### *Fig. S9*

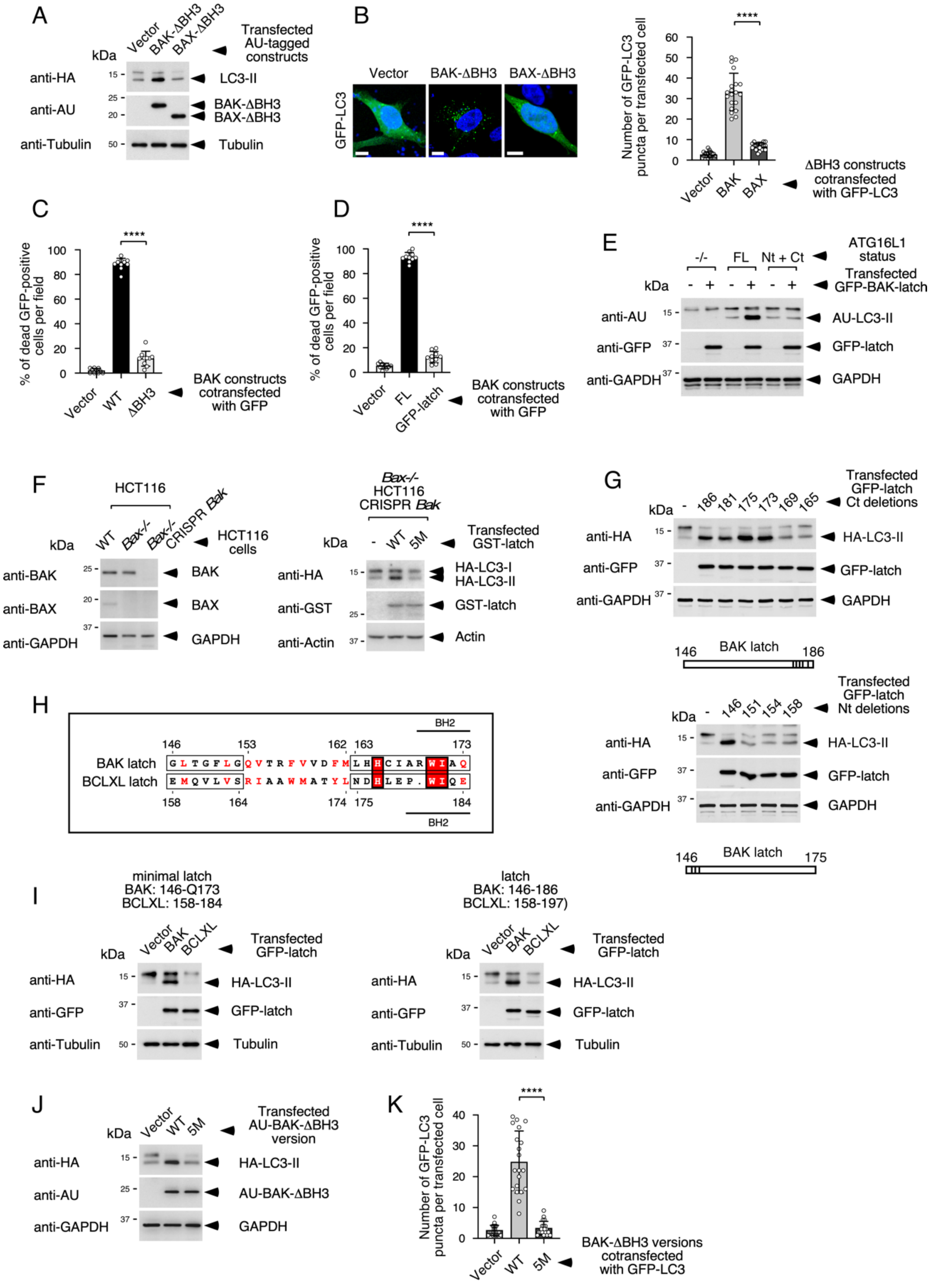

A
Vector
BAK-ΔBH3
BAX-ΔBH3
Transfected AU-tagged constructs
kDa
anti-HA
LC3-II
anti-AU
BAK-ΔBH3
BAX-ΔBH3
anti-Tubulin
Tubulin
B
Vector
BAK-ΔBH3
BAX-ΔBH3
GFP-LC3
Number of GFP-LC3 puncta per transfected cell
****
ΔBH3 constructs cotransfected with GFP-LC3
C
% of dead GFP-positive cells per field
****
Vector
WT
ΔBH3
BAK constructs cotransfected with GFP
D
FL
GFP-latch
E
ATG16L1 status
-/-
FL
Nt + Ct
Transfected GFP-BAK-latch
AU-LC3-II
anti-GFP
GFP-latch
anti-GAPDH
GAPDH
F
HCT116
Bax-/-
Bax-/- CRISPR Bak
HCT116 cells
anti-BAK
BAK
anti-BAX
BAX
Bax-/- HCT116 CRISPR Bak
5M
Transfected GST-latch
HA-LC3-I
HA-LC3-II
anti-GST
GST-latch
anti-Actin
Actin
G
Transfected GFP-latch Ct deletions
HA-LC3-II
BAK latch
Transfected GFP-latch Nt deletions
H
BH2
BAK latch
BCLXL latch
I
minimal latch
BAK: 146-Q173
BCLXL: 158-184
BAK
BCLXL
Transfected GFP-latch
latch
BAK: 146-186
BCLXL: 158-197)
J
Transfected AU-BAK-ΔBH3 version
AU-BAK-ΔBH3
K
BAK-ΔBH3 versions cotransfected with GFP-LC3

**Suppl. Fig. 9. BAK induces unconventional autophagy through the latch domain. (A-D)** BAK-ΔBH3 induces LC3 activation but not cell death. **(A)** HEK-293T cells were transfected with BAK- or BAX-ΔBH3 and HA-LC3, and lysed 36 h later for Western blot. **(B)** Hela cells were transfected with BAK- or BAX-ΔBH3 and GFP-LC3, and processed 36 h later for microscopy. Shown are representative confocal pictures (left) and quantification of GFP-LC3-positive puncta per cell (right; n = 20 cells). **(C, D)** HEK-293T cells were transfected with the indicated BAK constructs and GFP, and scored for cell death in the GFP-positive compartment 36 h later (n = 10 fields with at least 20 GFP-positive cells per field). **(E)** The BAK latch domain does not induce LC3 lipidation in HCT116 cells expressing separated Nt and Ct domains of ATG16L1. Cells were transfected with GFP-BAK latch (146-186) and AU-LC3, and lysed 36 h later for Western blot. **(F)** The BAK latch domain induces LC3 lipidation in the absence of endogenous BAK and BAX. *Bax-/-* HCT116 cells were targeted for BAK depletion using the CRISPR/Cas9 system (left), transfected with the shown GST-latch chimera and HA-LC3 and lysed 36 h post-transfection for Western blot (right). **(G-K)** Structure-function analysis of the BAK latch domain. **(G)** HEK-293T cells were transfected with the shown C-terminal (top) or N-terminal (bottom) latch deletions fused to GFP and HA-LC3, and lysed 36 h later for Western blot. **(H)** Sequence of the minimally active BAK latch domain aligned with the same region of BCLXL. Functionally conserved residues are in red font. Red boxes indicate identical residues. Boxes and numbers indicate the limits of the BAK/BCLXL latch chimeras analyzed in Fig. 4E. **(I)** The BCLXL latch domain fused to BAK TM does not cause LC3 lipidation. HEK-293T cells were transfected with the minimally active (left) or full-length (right) BAK and BCLXL latch domains (fused to GFP and the BAK TM) and HA-LC3, and lysed 36 h later for Western blotting. **(J, K)** BAK-ΔBH3-5M does not activate LC3. **(J)** HEK-293T cells were transfected with WT or 5M BAK-ΔBH3 and HA-LC3, and lysed 36 h later for Western blot. **(K)** Hela cells were transfected with WT or 5M BAK-ΔBH3 and GFP-LC3, and fixed 36 h later for microscopy to score the number of GFP-LC3-positive puncta per cell (n = 20 cells). All graphs in this figure show mean values -/+ s.d. (****$P$<0.0001 Student's $t$-test).

***Fig. S10***

A

| ACCESSION | DESCRIPTION | WT | 5M | DIFF. |
|---|---|---|---|---|
| F2Z2X4 | Exportin-4 - [F2Z2X4_HUMAN] | 21 | 1 | 20 |
| O14980 | Exportin-1 - [XPO1_HUMAN] | 33 | 17 | 16 |
| O00410 | Importin-5 - [IPO5_HUMAN] | 19 | 3 | 16 |
| J3KPX7 | Prohibitin-2 - [J3KPX7_HUMAN] | 13 | 1 | 12 |
| P55060 | Isoform 3 of Exportin-2 - [XPO2_HUMAN] | 28 | 17 | 11 |
| Q14974 | Importin subunit beta-1 - [IMB1_HUMAN] | 13 | 3 | 10 |
| P35232 | Prohibitin - [PHB_HUMAN] | 18 | 9 | 9 |
| O95373 | Importin-7 - [IPO7_HUMAN] | 16 | 8 | 8 |
| Q92973 | Isoform 2 of Transportin-1 - [TNPO1_HUMAN] | 14 | 6 | 8 |
| P21796 | Voltage-dependent anion-selective channel protein 1 - [VDAC1_HUMAN] | 9 | 2 | 7 |
| O14787 | Isoform 2 of Transportin-2 - [TNPO2_HUMAN] | 8 | 1 | 7 |
| Q9Y5L0 | Isoform 3 of Transportin-3 - [TNPO3_HUMAN] | 7 | 0 | 7 |
| P27105 | Erythrocyte band 7 integral membrane protein - [STOM_HUMAN] | 7 | 0 | 7 |
| O15397 | Importin-8 - [IPO8_HUMAN] | 8 | 2 | 6 |
| Q9UJZ1 | Stomatin-like protein 2, mitochondrial - [STML2_HUMAN] | 10 | 6 | 4 |
| Q9HAV4 | Exportin-5 - [XPO5_HUMAN] | 9 | 5 | 4 |
| Q96QU8 | Isoform 2 of Exportin-6 - [XPO6_HUMAN] | 5 | 1 | 4 |
| O43592 | Exportin-T - [XPOT_HUMAN] | 5 | 1 | 4 |

B

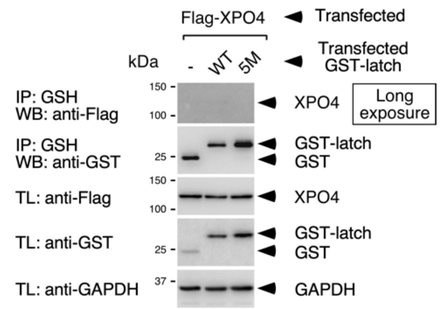

C

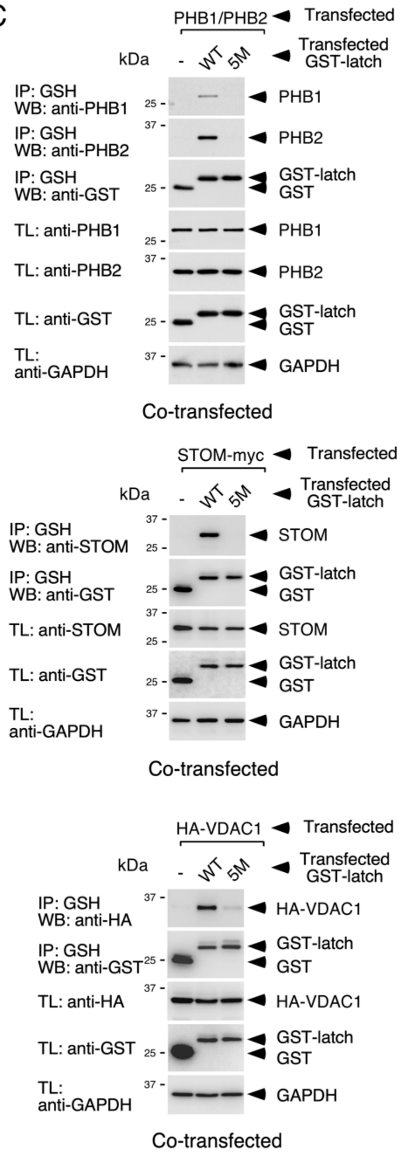

D

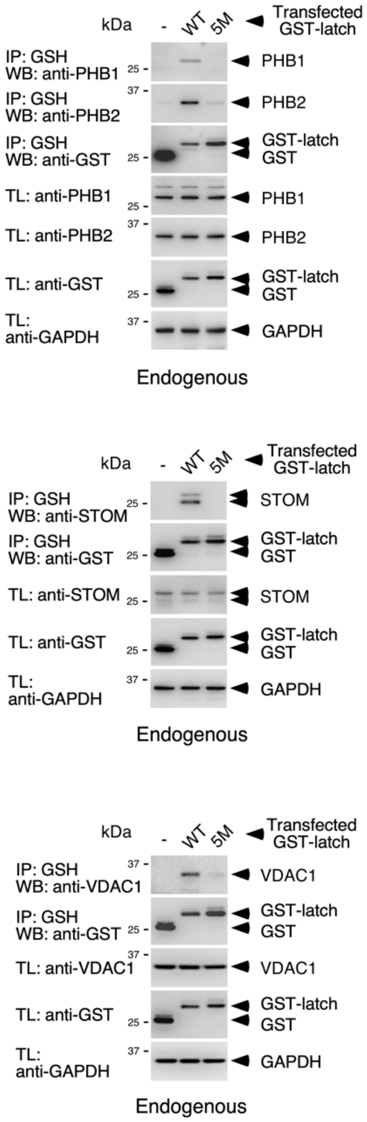

**Suppl. Fig. 10. The BAK latch domain specifically binds PHBs, STOM and VDAC1.** **(A)** Short list of selected proteins identified by proteomics as potential latch interactors. The list includes proteins over-represented by at least 4 peptides in the latch-WT sample over the 5M version and belonging to the Importin/Exportin and Prohibitin families. VDAC1 was selected since its family member VDAC2 was described as a BAK interactor. The number of peptides identified by mass spectrometry in the WT and 5M samples and the difference in these numbers (DIFF) are indicated. Prohibitin-family members and VDAC1 are highlighted. **(B)** EXPORTIN-4 does not coprecipitate with the latch domain. HEK-293T cells were transfected with the indicated versions of the BAK latch domain tagged with GST along with Flag-tagged EXPORTIN-4 (XPO4) and lysed 36 h post-transfection. The resulting extracts were incubated with GSH-agarose beads and the precipitates subjected to Western blot (IPs). Expression levels of all contenders in total protein lysates are shown on the bottom panels (TLs). **(C, D)** The BAK latch domain co-precipitates with co-transfected **(C)** and endogenous **(D)** PHBs, STOM, and VDAC1. HEK-293T cells were transfected with the indicated versions of the BAK latch domain tagged with GST in the presence **(C)** or absence **(D)** of vectors expressing PHB1 and PHB2 (top), STOM-myc (medium) or HA-VDAC (bottom), and lysed 36 h post-transfection. The resulting extracts were incubated with GSH-agarose beads and the precipitates subjected to Western blot (IPs). Expression levels of all contenders in total protein lysates are shown on the bottom panels (TLs).

***Fig. S11***

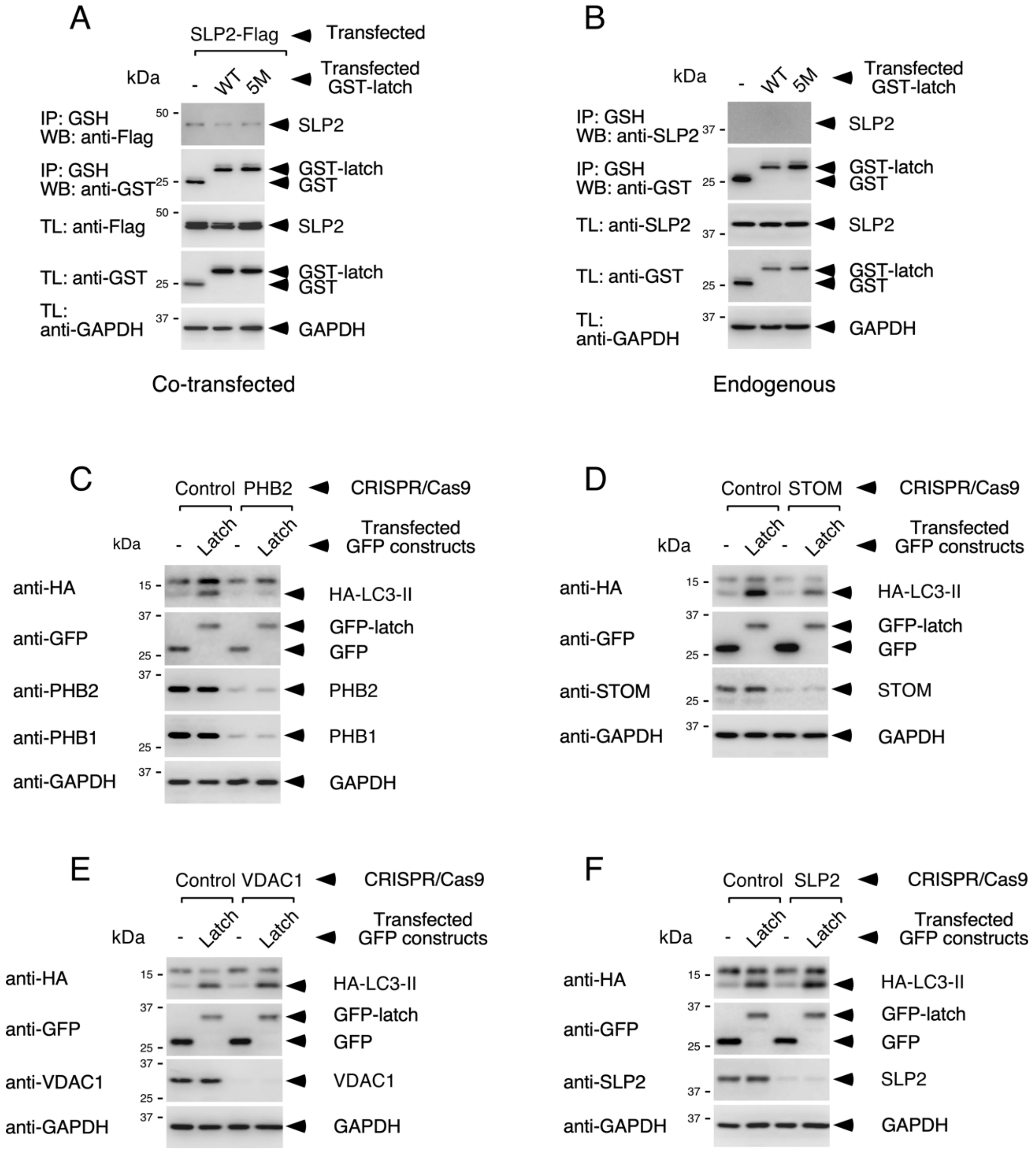
A
SLP2-Flag
Transfected
kDa
-
WT
5M
Transfected GST-latch
IP: GSH WB: anti-Flag
50
SLP2
IP: GSH WB: anti-GST
25
GST-latch
GST
TL: anti-Flag
50
SLP2
TL: anti-GST
25
GST-latch
GST
TL: anti-GAPDH
37
GAPDH
Co-transfected
B
kDa
-
WT
5M
Transfected GST-latch
IP: GSH WB: anti-SLP2
37
SLP2
IP: GSH WB: anti-GST
25
GST-latch
GST
TL: anti-SLP2
37
SLP2
TL: anti-GST
25
GST-latch
GST
TL: anti-GAPDH
37
GAPDH
Endogenous
C
Control
PHB2
CRISPR/Cas9
kDa
-
Latch
-
Latch
Transfected GFP constructs
anti-HA
15
HA-LC3-II
anti-GFP
37
25
GFP-latch
GFP
anti-PHB2
37
PHB2
anti-PHB1
25
PHB1
anti-GAPDH
37
GAPDH
D
Control
STOM
CRISPR/Cas9
kDa
-
Latch
-
Latch
Transfected GFP constructs
anti-HA
15
HA-LC3-II
anti-GFP
37
25
GFP-latch
GFP
anti-STOM
25
STOM
anti-GAPDH
37
GAPDH
E
Control
VDAC1
CRISPR/Cas9
kDa
-
Latch
-
Latch
Transfected GFP constructs
anti-HA
15
HA-LC3-II
anti-GFP
37
25
GFP-latch
GFP
anti-VDAC1
37
25
VDAC1
anti-GAPDH
37
GAPDH
F
Control
SLP2
CRISPR/Cas9
kDa
-
Latch
-
Latch
Transfected GFP constructs
anti-HA
15
HA-LC3-II
anti-GFP
37
25
GFP-latch
GFP
anti-SLP2
37
SLP2
anti-GAPDH
37
GAPDH

**Suppl. Fig. 11. The BAK latch domain does not interact with SLP2 and requires PHBs and STOM to induce LC3 activation. (A, B)** The BAK latch domain does not co-precipitate with co-transfected **(A)** or endogenous **(B)** SLP2. HEK-293T cells were transfected with the indicated versions of the BAK latch domain tagged with GST in the presence **(A)** or absence **(B)** of a vector expressing SLP2, and lysed 36 h post-transfection. The resulting extracts were incubated with GSH-agarose beads and the precipitates subjected to Western blot (IPs). Expression levels of all contenders in total protein lysates are shown on the bottom panels (TLs). **(C-F)** PHBs and STOM, but not VDAC1 or SLP2, are required for LC3 activation induced by the BAK latch domain. HEK-293T cells were transduced with CRISPR/Cas9 constructs targeting PHB2 **(C)**, STOM **(D)**, VDAC **(E)** or SLP2 **(F)** and after 58 h (for PHB2) or a week (for STOM, VDAC and SLP2), transfected with GFP or a GFP-latch fusion construct (as indicated). Cells were lysed 24 h post-transfection and subjected to Western blot.

### *Fig. S12*

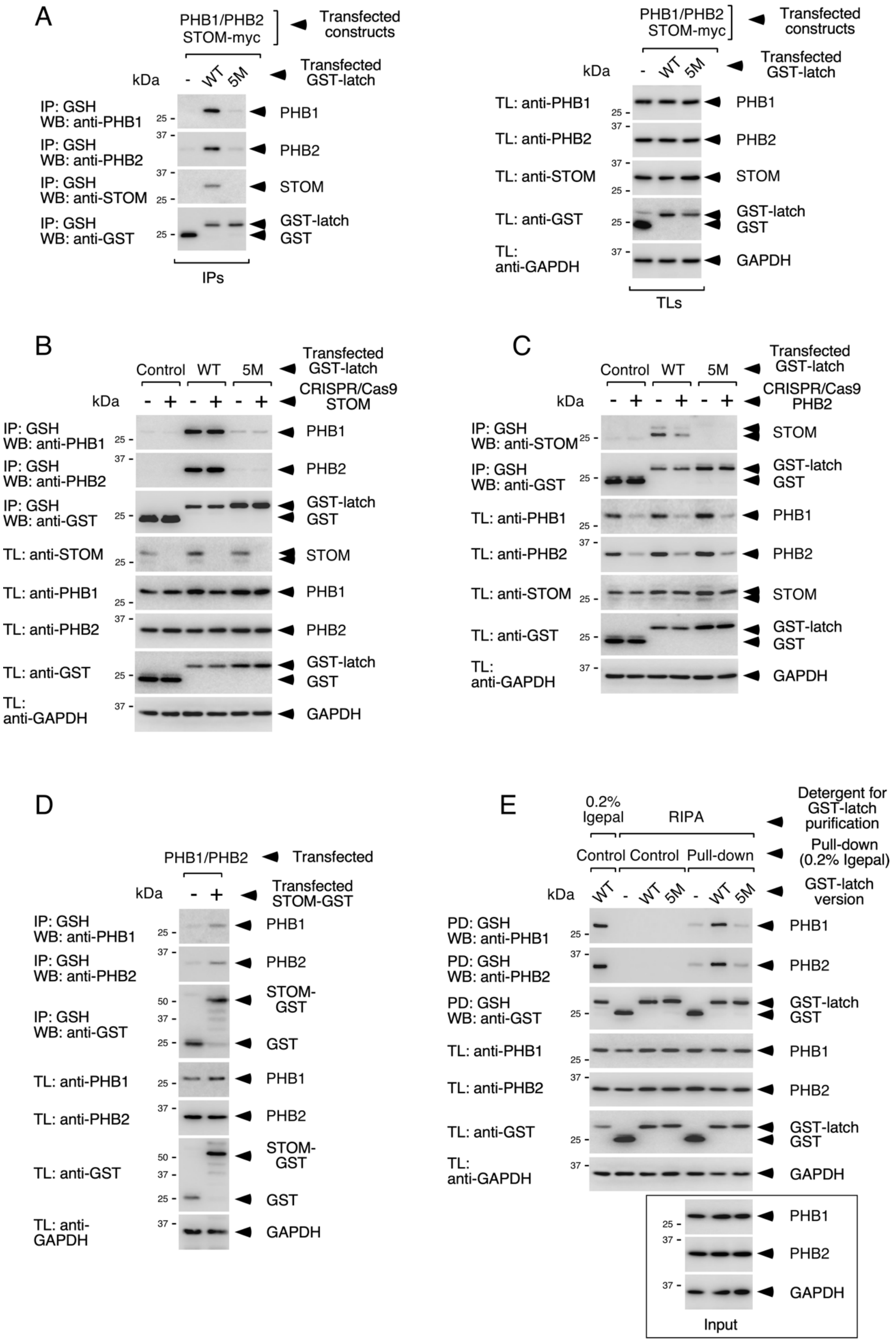
A
PHB1/PHB2
STOM-myc
Transfected constructs
Transfected GST-latch
kDa
-
WT
5M
IP: GSH
WB: anti-PHB1
WB: anti-PHB2
WB: anti-STOM
WB: anti-GST
PHB1
PHB2
STOM
GST-latch
GST
IPs
TL: anti-PHB1
TL: anti-PHB2
TL: anti-STOM
TL: anti-GST
TL: anti-GAPDH
GAPDH
TLs
B
Control
CRISPR/Cas9 STOM
C
CRISPR/Cas9 PHB2
D
PHB1/PHB2
Transfected
Transfected STOM-GST
STOM-GST
E
0.2% Igepal
RIPA
Detergent for GST-latch purification
Control
Pull-down
Pull-down (0.2% Igepal)
GST-latch version
PD: GSH
Input
25
37
50

**Suppl. Fig. 12. PHBs facilitate binding of STOM to the BAK latch domain (A)** Simultaneous co-precipitation of PHBs and STOM with the latch domain. HEK-293T cells were transfected with the indicated versions of the BAK latch domain tagged with GST along with PHBs and STOM-myc and lysed 36 h post-transfection. Lysates were incubated with GSH-agarose beads and the precipitates subjected to Western blot (left panel, IPs). Expression levels of all contenders in total protein lysates are shown on the right panel (TLs). **(B, C)** PHB depletion inhibits binding of endogenous STOM to the latch domain. HEK-293T cells were transduced with the shown CRISPR/Cas9 constructs to reduce STOM **(B)** or PHB **(C)** expression. Cells were then transfected with the indicated GST-latch chimeras and lysed 36 h post-transfection for GST precipitation and Western blot (IPs, as in **A**). Expression levels of all proteins in total lysates are shown on the bottom panels (TLs). **(D)** PHBs bind STOM-GST. HEK-293T cells were transfected with GST or a STOM-GST fusion protein along with PHB1 and PHB2 and lysed 36 h post-transfection. Lysates were incubated with GSH-agarose beads and the precipitates subjected to Western blot (IPs). Expression levels of all contenders in total protein lysates are shown on the bottom panels (TLs). **(E)** The wild-type BAK latch domain, but not the 5M mutant, binds PHBs in a pull-down assay. HEK-293T cells were transfected with plasmids expressing the indicated GST-latch versions and lysed 36 h post-transfection for GST purification in the shown detergent conditions. Beads containing GST chimera purified in harsh conditions (RIPA buffer) were used to pull-down transfected PHBs from total lysates under soft conditions (0.2% Igepal), and subjected to Western blotting along with control beads. The figure shows that beads isolated using RIPA buffer are devoid of endogenous PHBs (control lanes 2-4) compared to purification in 0.2% Igepal (lane 1), and are able to pull-down PHBs from total lysates overexpressing PHBs (input panels at the bottom) under soft conditions (Igepal 0.2%; pull-down lanes 5-7).

### *Fig. S13*

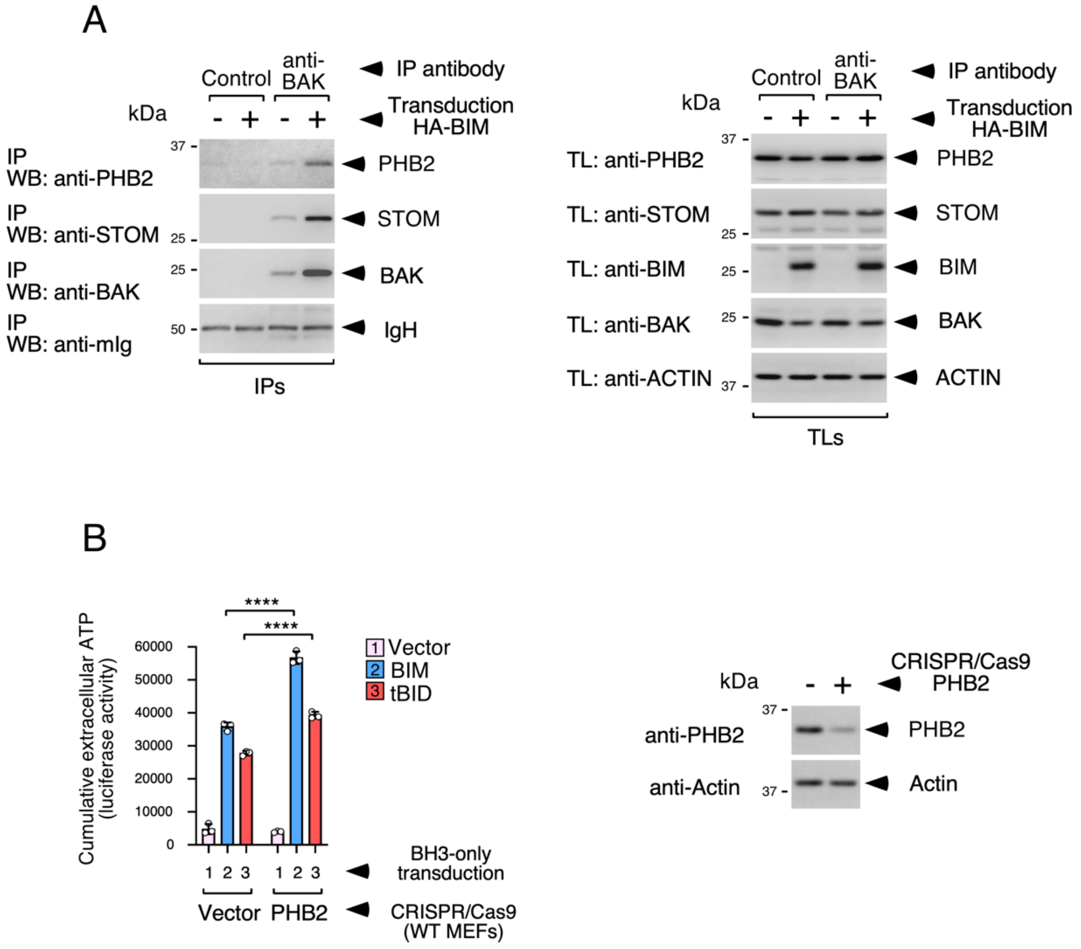
A
Control
anti-BAK
IP antibody
Transduction HA-BIM
kDa
- + - +
IP WB: anti-PHB2
IP WB: anti-STOM
IP WB: anti-BAK
IP WB: anti-mIg
37
25
25
50
PHB2
STOM
BAK
IgH
IPs
TL: anti-PHB2
TL: anti-STOM
TL: anti-BIM
TL: anti-BAK
TL: anti-ACTIN
PHB2
STOM
BIM
BAK
ACTIN
TLs
B
Cumulative extracellular ATP (luciferase activity)
60000
50000
40000
30000
20000
10000
0
****
****
1 Vector
2 BIM
3 tBID
1 2 3 1 2 3
Vector PHB2
BH3-only transduction
CRISPR/Cas9 (WT MEFs)
CRISPR/Cas9 PHB2
anti-PHB2
anti-Actin
PHB2
Actin

**Suppl. Fig. 13. PHBs and STOM assemble into a BAK-nucleated complex in response to BIM that inhibits ATP release during apoptosis. (A)** BAK activated by BIM binds PHB2 and STOM. DKO MEFs depleted of CASP3 and CASP9 via CRISPR/Cas9 and expressing human BAK (and also human PHBs and STOM for increased detectability), were transduced with HA-BIM and, 15 h later, crosslinked with 1% paraformaldehyde and lysed for immunoprecipitation with an anti-BAK antibody that binds the exposed Nt domain of BAK (G317.2) or a control immunoglobulin (control) (IPs, left panel). Expression levels of all molecules in total protein lysates are shown on the right panel (TLs). **(B)** PHB depletion increases ATP release during apoptosis in wild-type MEFs. Wild-type MEFs were transduced with the shown CRISPR/Cas9 constructs and, 40 h later, transduced to express BH3-only molecules for 22 h. The levels of extracellular ATP were measured at 14 h and 22 h. The graph shows cumulative ATP release data resulting from the sum of the luciferase activity units obtained at both time points. Shown are mean values -/+ s.d. from triplicate experimental points (n = 3; ****$P$<0.0001 Student's $t$-test). Separate control points were lysed 48 h post lentiviral CRISPR/Cas9 transduction to determine PHB2 depletion by Western blot (right panel).

***Fig. S14***

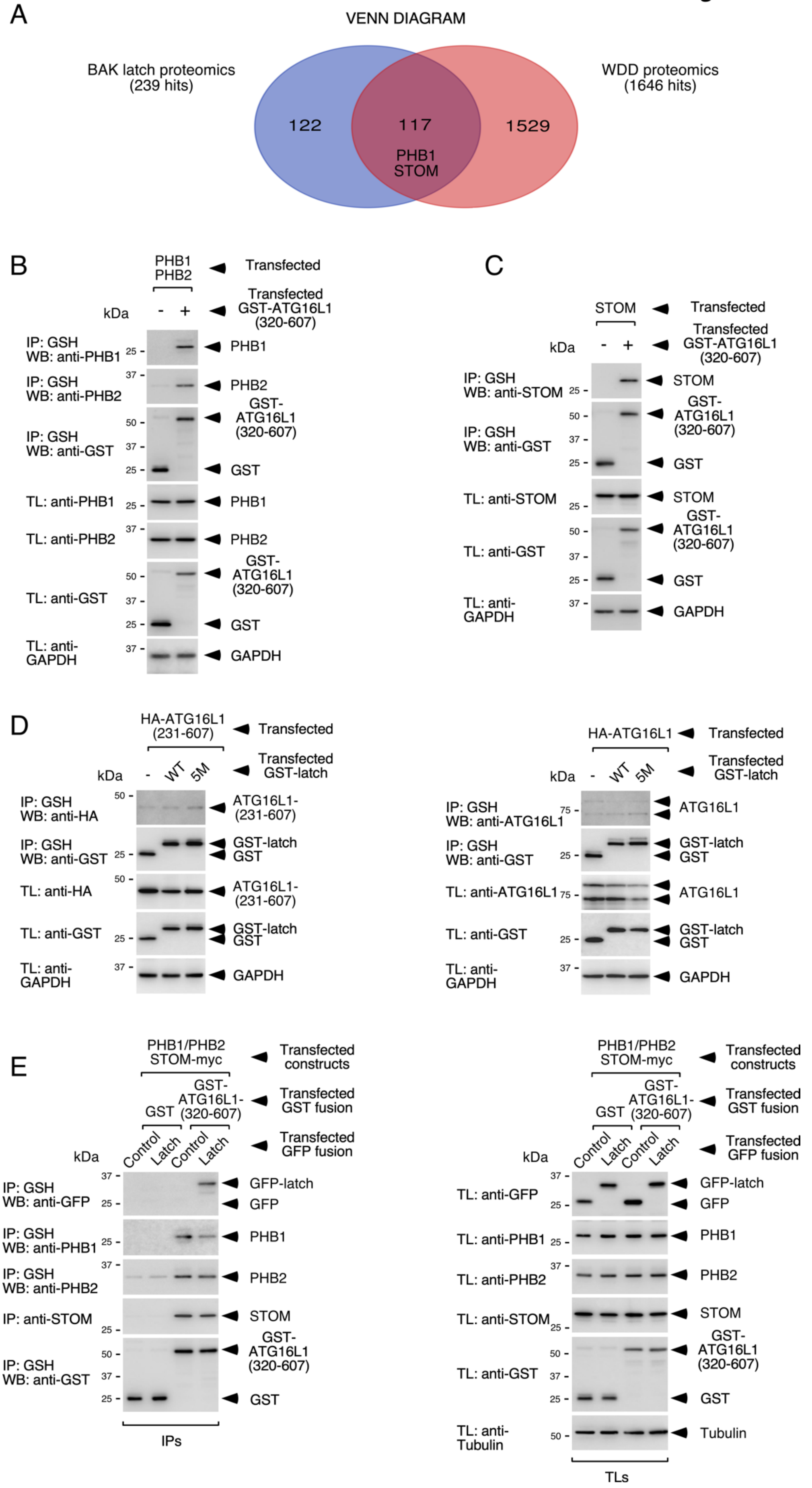
A
VENN DIAGRAM
BAK latch proteomics
(239 hits)
WDD proteomics
(1646 hits)
122
117
1529
PHB1
STOM
B
PHB1
PHB2
Transfected
Transfected
GST-ATG16L1
(320-607)
kDa
IP: GSH
WB: anti-PHB1
IP: GSH
WB: anti-PHB2
IP: GSH
WB: anti-GST
TL: anti-PHB1
TL: anti-PHB2
TL: anti-GST
TL: anti-
GAPDH
PHB1
PHB2
GST-
ATG16L1
(320-607)
GST
GAPDH
C
STOM
Transfected
Transfected
GST-ATG16L1
(320-607)
kDa
IP: GSH
WB: anti-STOM
IP: GSH
WB: anti-GST
TL: anti-STOM
TL: anti-GST
TL: anti-
GAPDH
STOM
GST-
ATG16L1
(320-607)
GST
GAPDH
D
HA-ATG16L1
(231-607)
Transfected
Transfected
GST-latch
kDa
WT
5M
IP: GSH
WB: anti-HA
IP: GSH
WB: anti-GST
TL: anti-HA
TL: anti-GST
TL: anti-
GAPDH
ATG16L1-
(231-607)
GST-latch
GST
GAPDH
HA-ATG16L1
IP: GSH
WB: anti-ATG16L1
TL: anti-ATG16L1
ATG16L1
E
PHB1/PHB2
STOM-myc
Transfected
constructs
GST
GST-
ATG16L1-
(320-607)
Transfected
GST fusion
Control
Latch
Transfected
GFP fusion
IP: GSH
WB: anti-GFP
IP: GSH
WB: anti-PHB1
IP: GSH
WB: anti-PHB2
IP: anti-STOM
IP: GSH
WB: anti-GST
GFP-latch
GFP
PHB1
PHB2
STOM
IPs
TL: anti-GFP
TL: anti-PHB1
TL: anti-PHB2
TL: anti-STOM
TL: anti-GST
TL: anti-
Tubulin
Tubulin
TLs

**Suppl. Fig. 14. PHBs and STOM bind the WD40 domain of ATG16L1. (A)** Venn diagram of the BAK latch proteomics results and the list of possible interactors of the ATG16L1 WD40 domain previously identified. The pool of overlapping proteins includes PHB1 and STOM. **(B, C)** PHBs and STOM co-precipitate with GST-WD40 (320-607). HEK-293T cells were transfected with the indicated PHB **(B)** or STOM **(C)** constructs along with GST or GST-WD40 domain (320-607), lysed 36 h later and the resulting extracts incubated with GSH-agarose beads for precipitation and Western blot (IPs). Expression levels of all contenders in total lysates are also shown (TLs, bottom panels). **(D)** The BAK latch domain does not bind ATG16L1. HEK-293T cells were transfected with the shown GST-latch fusions along with HA-ATG16L1-(231-607) (left) or full-length ATG16L1 (right) and lysed 36 h post-transfection for GST precipitation and Western blot (IPs). The anti-HA (left) and anti-ATG16L1 (right) IP panels were overexposed. Expression levels of all proteins in total lysates are also shown (TLs). **(E)** PHBs and STOM facilitate co-precipitation of the ATG16L1 WD40 domain with GFP-latch. HEK-293T cells were transfected with GST or GST-ATG16L1 (320-607) along with PHBs, STOM-myc and the indicated GFP-latch constructs, and lysed 36 h post-transfection for GST precipitation and Western blot (IPs, left panel). Expression levels of all proteins in total lysates are shown on the right panel (TLs).

***Fig. S15***

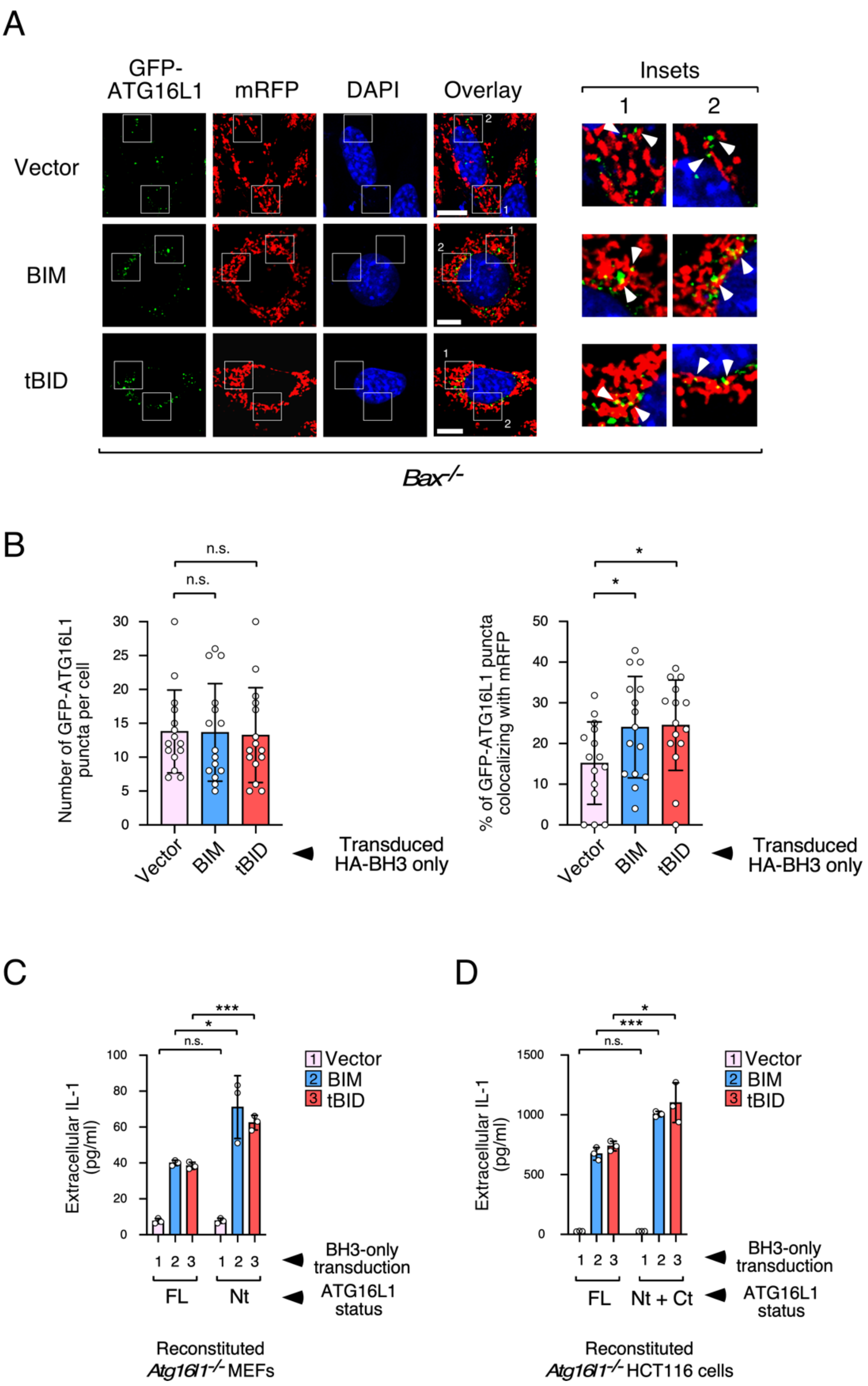
A
GFP-ATG16L1
mRFP
DAPI
Overlay
Insets
1
2
Vector
BIM
tBID
Bax-/-
B
Number of GFP-ATG16L1 puncta per cell
n.s.
n.s.
Vector
BIM
tBID
Transduced HA-BH3 only
% of GFP-ATG16L1 puncta colocalizing with mRFP
*
*
Transduced HA-BH3 only
C
Extracellular IL-1 (pg/ml)
n.s.
*
***
1 Vector
2 BIM
3 tBID
BH3-only transduction
FL
Nt
ATG16L1 status
Reconstituted Atg16l1-/- MEFs
D
Extracellular IL-1 (pg/ml)
n.s.
***
*
1 Vector
2 BIM
3 tBID
BH3-only transduction
FL
Nt + Ct
ATG16L1 status
Reconstituted Atg16l1-/- HCT116 cells

**Suppl. Fig. 15. ATG16L1 is recruited to mitochondria and its WD40 domain is required for inhibition of IL-1β release by co-cultured BMDMs. (A)** Recruitment of ATG16L1 to mitochondria caused by BH3-only proteins. *Bak-/-* MEFs expressing GFP-ATG16L1 and mitochondrial RFP (mRFP) were transduced with BH3-only molecules, treated with zVAD.fmk 7 h later and fixed for microscopy 17 h post-transduction. Shown are representative confocal pictures. White arrows indicate GFP-ATG16L1 vesicles that do (BIM, tBID) or do not (Vector) closely appose to mRFP-labelled mitochondria to generate a yellow signal. **(B)** Quantification of the phenotype shown in **A**. Graphs display mean values -/+ s.d. of the number of GFP-ATG16L1 puncta per cell (left; n = 15 cells) and the percentage of GFP-ATG16L1 puncta that colocalize with mRFP producing a yellow signal (right; n = 15 cells), n.s. $P>0.05$, *$P<0.05$ Student's *t*-test. **(C, D)** The WD40 domain in the dying cells suppresses IL-1β release by co-cultured BMDMs. *Atg16l1-/-* MEFs reconstituted with ATG16L1 FL or Nt (1-299) **(C)** or *Atg16l1-/-* HCT116 cells expressing ATG16L1 FL or Nt + Ct (300-607) **(D)** were transduced with BH3-only molecules and, 7 h later, treated with 3 μM of ARL67156. Apoptotic bodies were collected 20 h **(C)** or 24 h **(D)** post-transduction and tested on pre-activated BMDMs for IL-1β secretion by ELISA. Graph shows mean values of IL-1β concentration -/+ s.d. from triplicate experimental points (n = 3; n.s. $P>0.05$, *$P<0.05$, ***$P<0.001$ Student's *t*-test).

### Supplementary Tables

### Suppl. Table 1 (oligonucleotides).

| | |
|---|---|
| **BH3-only and VDAC (PCR)** | |
| Hind3-Kozak-Nco1-BIM-fw | gggcccaagcttgccaccatggcaaagcaaccttctgatg |
| Not1-Stop-BIM-rev | cccggggcggccgctttaatgcattctccacaccaggcgg |
| Hind3-Kozak-Nco1-BAD-fw | gggcccaagcttgccaccatggtccagatcccagagtttgagcc |
| Not1-Stop-BAD-rev | cccggggcggccgctttactgggagggggcggagcttccc |
| Hind3-Kozak-Nco1-tBID-fw | gggcccaagcttgccaccatgggcaaccgcagcagccactccc |
| Not1-Stop-tBID-rev | cccggggcggccgctttagtccatcccatttctggctaag |
| Pci1-PUMA-fw | gggcccacatgtccggcatggcccgcgcacgccaggagggc |
| Not1-Stop-PUMA-rev | cccggggcggccgctttaattgggctccatctcgggggctc |
| Pci1-BIK-fw | gggcccacatgtctgaagtaagacccctctc |
| Not1-Stop-BIK-rev | cccggggcggccgctttacttgagcagcaggtgcaggcccc |
| Pci1-NOXA-fw | gggcccacatgtccggcatgcctgggaagaaggcgcgcaag |
| Not1-Stop-NOXA-rev | cccggggcggccgctttaggttcctgagcagaagagtttgg |
| Pci1-bNIP3L-fw | gggcccacatgtcgtcccacctagtcgagccgcc |
| Not1-Stop-bNIP3L-rev | cccggggcggccgctttagtaggtgctggcagagggtgtgctca |
| Pci1-bNIP3-fw | gggcccacatgtcgcagaacggagcgcccgggatg |
| Not1-Stop-bNIP3-rev | cccggggcggccgctttaaaaggtgctggtggaggttgtcagac |
| Nco1-VDAC1-fw | gggcccccatggctgtgccacccacgtatgc |
| Not1-Stop-VDAC1-rev | cccggggcggccgctttatgcttgaaattccagtcctagacc |
| **BH3-only and BAK BH3 domain mutants (mutagenesis)** | |
| BIMEL-L152E-TOP | atggatcgcccaagaggagcggcgtatcggagacg |
| BIMEL-L152E-BOTT | cgtctccgatacgccgctcctcttgggcgatccat |
| BID-L90E-TOP | gaatattgccaggcacgaggcccaggtcggggacag |
| BID-L90E-BOTT | ctgtccccgacctgggcctcgtgcctggcaatattc |
| PUMA-L141E-TOP | ggagatcggggcccaggagcggcggatggcggacg |
| PUMA-L141E-BOTT | cgtccgccatccgccgctcctgggccccgatctcc |
| BAK-L78E-TOP | gcaggtgggacggcaggaggccatcatcggggacg |
| BAK-L78E-BOTT | cgtccccgatgatggcctcctgccgtcccacctgc |
| **BAK deletions (PCR)** | |
| Pci1-BAK-fw | gggcccacatgtccggcggcatggcttcggggcaaggcccaggt |
| Not1-Stop-BAK-rev | cccggggcggccgctttatgatttgaagaatcttcgtaccac |
| Pci1-BAK-G146-fw | gggcccacatgtccggcctgactggcttcctaggccaggtgacc |
| Mlu1-BAK-M71-rev | cccgggacgcgtcatggtgctgctaggttgcagagg |
| Pci1-BAK-Y89-fw | gggcccacatgtcctatgactcagagttccagaccatgttg |
| Mlu1-BAK-G146-rev | cccgggacgcgtgccatgctggtagacgtgtagggcc |
| EcoR1-Mlu1-BAK-TM-fw | gggcccgaattcacgcgtcccatcctgaacgtgctggtggttctgg |
| EcoR1-BAK-deltaBH3-rev | cccggggaattccatggtgctgctaggttgcaga |
| EcoR1-BAK-deltaBH3-fw | gggcccgaattctatgactcagagttccagacca |
| **BAX deletions (PCR)** | |
| Nco1-BAX-fw | gggcccccatggacgggtccggggagcagc |
| Not1-Stop-BAX-rev | cccggggcggccgctttagcccatcttcttccagatggtg |
| Pci1-BAX-T127-fw | gggcccacatgtccaccaaggtgccggaactgatcagaac |
| Mlu1-BAX-T56-rev | cccgggacgcgtggtggacgcatcctgaggcaccggg |
| Nco1-BAX-M74-fw | gggcccccatgggcatggagctgcagaggatgattgccgccgtg |
| Mlu1-BAX-T127-rev | cccgggacgcgtggtgcacagggccttgagcaccagtttgctggc |
| EcoR1-Mlu1-BAX-TM-fw | gggcccgaattcacgcgtcagaccgtgaccatctttgtggcgggag |
| EcoR1-BAX-deltaBH3-rev | cccggggaattccttcttggtggacgcatcctgaggcac |
| EcoR1-BAX-deltaBH3-fw | gggcccgaattcatggagctgcagaggatgattgccgc |

| | |
|---|---|
| **BAK latch (146-186) PCR** | |
| Pci1-BAK-G146-fw | gggcccacatgtccggcctgactggcttcctaggccaggtgacc |
| Mlu1-BAK-G186-rev | cccgggacgcgtaccattgcccaagttcagggctgccaccc |
| **BAK latch (146-173) PCR** | |
| Pci1-BAK-G146-fw | gggcccacatgtccggcctgactggcttcctaggccaggtgacc |
| Mlu1-BAK-Q173-rev | cccgggacgcgtctgtgcaatccaccgggcaatgcagtg |
| **BAX latch (127-170) PCR** | |
| Pci1-BAX-T127-fw | gggcccacatgtccaccaaggtgccggaactgatcagaac |
| Mlu1-BAX-W170-rev | cccgggacgcgtccacgtgggcgtcccaaagtaggagagg |
| **BCLXL latch (158-197) PCR** | |
| Pci1-BCLXL-E158-fw | gggcccacatgtccgagatgcaggtattggtgagtcggatc |
| Mlu1-BCLXL-N197-rev | cccgggacgcgtgttcccatagagttccacaaaagtatcc |
| **BCLXL latch (158-184) PCR** | |
| Pci1-BCLXL-E158-fw | gggcccacatgtccgagatgcaggtattggtgagtcggatc |
| Mlu1-BCLXL-E184-rev | cccgggacgcgtctcctggatccaaggctctaggtggtc |
| **Ct BAK latch deletions PCR** | |
| Pci1-BAK-G146-fw | gggcccacatgtccggcctgactggcttcctaggccaggtgacc |
| Mlu1-BAK-G186-rev | cccgggacgcgtaccattgcccaagttcagggctgccaccc |
| Mlu1-BAK-L181-rev | cccgggacgcgtcagggctgccacccagccacccctctg |
| Mlu1-BAK-G175-rev | cccgggacgcgtacccctctgtgcaatccaccgggcaatg |
| Mlu1-BAK-Q173-rev | cccgggacgcgtctgtgcaatccaccgggcaatgcagtg |
| Mlu1-BAK-R169-rev | cccgggacgcgtccgggcaatgcagtgatgcagcatgaag |
| Mlu1-BAK-H165-rev | cccgggacgcgtgtgatgcagcatgaagtcgaccacgaagc |
| **Nt BAK latch deletions PCR** | |
| Mlu1-BAK-G175-rev | cccgggacgcgtacccctctgtgcaatccaccgggcaatg |
| Pci1-BAK-G146-fw | gggcccacatgtccggcctgactggcttcctaggccaggtgacc |
| Pci1-BAK-L151-fw | gggcccacatgtccctaggccaggtgacccgcttcgtggtc |
| Pci1-BAK-V154-fw | gggcccacatgtccgtgacccgcttcgtggtcgacttcatg |
| Pci1-BAK-V158-fw | gggcccacatgtccgtggtcgacttcatgctgcatcactgc |
| **BAK-BCLXL latch chimeric constructs (PCR)** | |
| **Chimera 2: BCLXL-latch (158-164): BAK-latch (153-173)** | |
| Pci1-BCLXL-(E158-S164)-BAK-fw | gggcccacatgtccgagatgcaggtgctggtgagccaggtgacccgcttcgtggtcgacttcatg |
| Mlu1-BAK-Q173-rev | cccgggacgcgtctgtgcaatccaccgggcaatgcagtg |
| **Chimera 3: BAK-latch (146-162): BCLXL-latch (175-184)** | |
| Pci1-BAK-G146-fw | gggcccacatgtccggcctgactggcttcctaggccaggtgacc |
| Mlu1-BCLXL-(N175-E184)-BAK-rev | cccgggacgcgtctcctggatccaaggctccaggtggtcgttcatgaagtcgaccacgaagcgggtcacctg |
| **Chimera 4: BCLXL-latch (158-174): BAK-latch (163-173)** | |
| Pci1-BCLXL-E158-fw | gggcccacatgtccgagatgcaggtattggtgagtcggatc |
| Mlu1-BAK-(N163-Q173)-BCLXL-rev | cccgggacgcgtctgtgcaatccaccgggcaatgcagtgatgcagcaggtaggtggccatccaagcggc |
| **BAK latch 5M (mutagenesis)** | |
| BAK-NDHLEP-TOP | gacccgcttcgtggtcgacttcatgaacgaccacctggagccccggtggattgcacagaggggtggct |
| BAK-NDHLEP-BOTT | agccacccctctgtgcaatccaccggggctccaggtggtcgttcatgaagtcgaccacgaagcgggtc |

| **sgRNA GUIDES FOR LENTICRISPR** | |
|---|---|
| mouse ATG16L1 #1 (36-55 plus)-TOP | caccgctggaagcgtcacatcgcgg |
| mouse ATG16L1 #1 (36-55 plus)-BOTT | aaacccgcgatgtgacgcttccagc |
| mouse ATG16L1 #2 (1073-1092 plus)-TOP | caccgtggaccgcagggtgaaactt |
| mouse ATG16L1 #2 (1073-1092 plus)-BOTT | aaacaagtttcaccctgcggtccac |
| human STOM (minus)-TOP | caccgaaatccatccgcaaggtcca |
| human STOM (minus)-BOTT | aaactggaccttgcggatggatttc |
| human VDAC1 (plus)-TOP | caccgccacccacgtatgccgatct |
| human VDAC1 (plus)-BOTTOM | aaacagatcggcatacgtgggtggc |
| human PHB2 (plus)-TOP | caccgatcttcttcaatcggatcgg |
| human PHB2 (plus)-BOTT | aaacccgatccgattgaagaagatc |
| mouse PHB2 (minus)-TOP | caccgcacggattcgcggacgccgt |
| mouse PHB2 (minus)-BOTT | aaacacggcgtccgcgaatccgtgc |
| mouse CASPASE-9 (minus)-TOP | caccgggcgcaccctgcatcgccgc |
| mouse CASPASE-9 (minus)-BOTT | aaacgcggcgatgcagggtgcgccc |
| mouse CASPASE-3 (plus)-TOP | caccgatctcgctctggtacggatg |
| mouse CASPASE-3 (plus)-BOTT | aaaccatccgtaccagagcgagatc |
| mouse STING (minus)-TOP | caccgcagtagtccaagttcgtgcg |
| mouse STING (minus)-BOTT | aaaccgcacgaacttggactactgc |
| human BAK (minus)-TOP | caccggaggtaaggtgaccatctct |
| human BAK (minus-BOTT | aaacagagatggtcaccttacctcc |
